# Tissue and regional expression patterns of dicistronic tRNA-mRNA transcripts in grapevine (*Vitis vinifera*) and their evolutionary co-appearance with vasculature in land plants

**DOI:** 10.1101/2020.04.13.039131

**Authors:** Pastor Jullian Fabres, Lakshay Anand, Na Sai, Stephen Pederson, Fei Zheng, Alexander A. Stewart, Benjamin Clements, Edwin R Lampugnani, James Breen, Matthew Gilliham, Penny Tricker, Carlos M. Rodríguez López, Rakesh David

**Author notes:** Corresponding author: Carlos M. Rodríguez López. Joint senior authorship (Carlos M. Rodríguez López, Rakesh David).

## Abstract

Transfer RNAs (tRNA) are crucial adaptor molecules between messenger RNA (mRNA) and amino acids. Recent evidence in plants suggests that dicistronic tRNA-like structures also act as mobile signals for mRNA transcripts to move between distant tissues. Co-transcription is not a common feature in the plant nuclear genome and, in the few cases where polycistronic transcripts have been found, they include non-coding RNA species such as small nucleolar RNAs and microRNAs. It is not known, however, the extent to which dicistronic transcripts of tRNA and mRNAs are expressed in field-grown plants, or the factors contributing to their expression. We analysed tRNA-mRNA dicistronic transcripts in the major horticultural crop grapevine (*Vitis vinifera*) using a novel pipeline developed to identify dicistronic transcripts from high-throughput RNA sequencing data. We identified dicistronic tRNA-mRNA in leaf and berry samples from 22 commercial vineyards. Of the 124 tRNA genes that were expressed in both tissues, 18 tRNA were expressed forming part of 19 dicistronic tRNA-mRNAs. The presence and abundance of dicistronic molecules was tissue and geographic sub-region specific. In leaves, the expression patterns of dicistronic tRNA-mRNAs significantly correlated with tRNA expression, suggesting that their transcriptional regulation might be linked. We also found evidence of syntenic genomic arrangements of tRNAs and protein coding genes between grapevine and *Arabidopsis thaliana*, and widespread prevalence of dicistronic tRNA-mRNA transcripts among vascular land plants but no evidence of these transcripts in nonvascular lineages. This suggests that the appearance of plant vasculature and tRNA-mRNA occurred concurrently during the evolution of land plants.

## Introduction

Polycistronic mRNAs are RNA molecules that contain two or more open reading frames (ORFs). These are usually found in viruses, bacteria, archaea, protozoans and invertebrates^1^. Polycistronic transcripts are synthesized when multiple genes forming an operon are co-expressed from a single promoter. These transcripts are then translated into protein from two or more translation initiation sites. This strategy has been described as an efficient mechanism to coordinate gene expression^1^. Although polycistronic transcripts are less common in plants, several chloroplast genes are organized in clusters and are co-transcribed in polycistronic primary transcripts and subsequently processed to form mature RNAs^2^, reflecting their prokaryotic ancestry^3^. The majority of nuclear-encoded genes in plants are monocistronic with a few exceptions, such as certain classes of polycistronic microRNAs (miRNAs)^4^ and small nucleolar RNAs (snoRNAs), which are organized in genomic clusters and are transcribed from a common promoter^5^. These precursor transcripts are processed to mature snoRNA and miRNA molecules. There are also a few reports of dicistronic transcripts encoding genes that are not functionally related to each other such as tRNAs-snoRNA, snoRNA-miRNAs and tRNA-mRNA in some plant species^5–7^; however, the molecular and physiological significance of co-transcription for many of these transcripts is still poorly understood.

Recent work in model plants, *Arabidopsis thaliana* and tobacco, has shed light on the function of dicistronic tRNAs-mRNAs^7^. Using transgenic lines, Zhang et al.^7^ demonstrated that tRNA-like structures (TLSs), when co-transcribed with mRNA transcripts, could act as mobility signals, triggering the systemic movement of the mRNA between roots and shoots. Notably, the mRNA components of the dicistronic transcripts were also shown to be translated into functional proteins. Endogenously produced tRNA-mRNA dicistronic transcripts have also been detected in *A. thaliana* suggesting that functional tRNA and tRNA-like structures could act as non-autonomous signals in plants able to deliver functional mRNAs to distantly located tissues. Beyond their canonical role in protein translation, tRNAs have been also demonstrated to function in other chemical transformations, for example, delivering amino acids during lipid modification and antibiotic biosynthesis^8^.

In grapevine (*Vitis vinifera*), the effect of growth environment on gene expression has been extensively studied^9,10^. Several studies have identified small non-coding RNAs (sRNAs) in grapevine that can influence development in response to environmental stimuli. Among these sRNAS, miRNAs respond to low temperature treatment^11^, application of exogenous gibberellin^12^ and viral infection^13^. In addition, studies have shown that miRNAs present tissue specificity in grapevine^14^. Bester et al.^15^ identified sRNA species in grapevine phloem. Notably, this study also showed the non-random manner in which tRNA-derived sRNAs originated^15^. A study looking at the effect of grafting in grapevine identified more than 3000 transcripts moving across graft junctions including transcripts for genes involved in the response to abiotic stress and signal transduction^16^. Zhang et al.^7^ confirmed that 11% of the mobile mRNA in *V. vinifera* from Yang et al. (2015)^16^ also had TLS motifs in their coding sequence or 3’ UTR.

We hypothesized that dicistronic tRNA-mRNA transcripts would be transcribed differentially between different grapevine tissues and in growing regions with different environments. As a first step towards identifying such transcripts, we present DiRT (**Di**cistronic **R**NA **T**ranscripts), a computational pipeline to detect dicistronic transcripts from short-read RNA-seq data that can be adapted for use in any organism. Using this pipeline, we analysed dicistronic tRNA-mRNA transcripts in commercial, field-grown grapevine and assessed the prevalence of these transcripts across the plant kingdom.

## Results

### RNA-sequencing of *Vitis vinifera* cv. Shiraz

To identify tRNA-mRNA dicistronic transcripts in *Vitis vinifera* cv. Shiraz, we performed RNA-seq of libraries from two different tissues, leaf and berry, collected at budburst) and veraison (onset of ripening) respectively, from 22 vineyards (own-rooted) from the Barossa wine growing region, South Australia, Australia (Figure 1). The wine region has been previously divided into different sub-regions according to the unique combinations of growing environments such as temperature, rainfall, soil type and elevation, which in turn contribute in differences in plant growth, berry composition and wine characteristics^17^. Sequencing reads were aligned to the 12x reference grapevine genome PN40024^18^ with an average mapping percentage of 90% for leaf and 87% for berry samples. We obtained an average of 23 million and 21 million paired-end (2×75 nucleotide) Illumina reads for each leaf and berry sample (three plants per sample, three samples per vineyard) (Supplemental Table S1).

**Figure 1.**
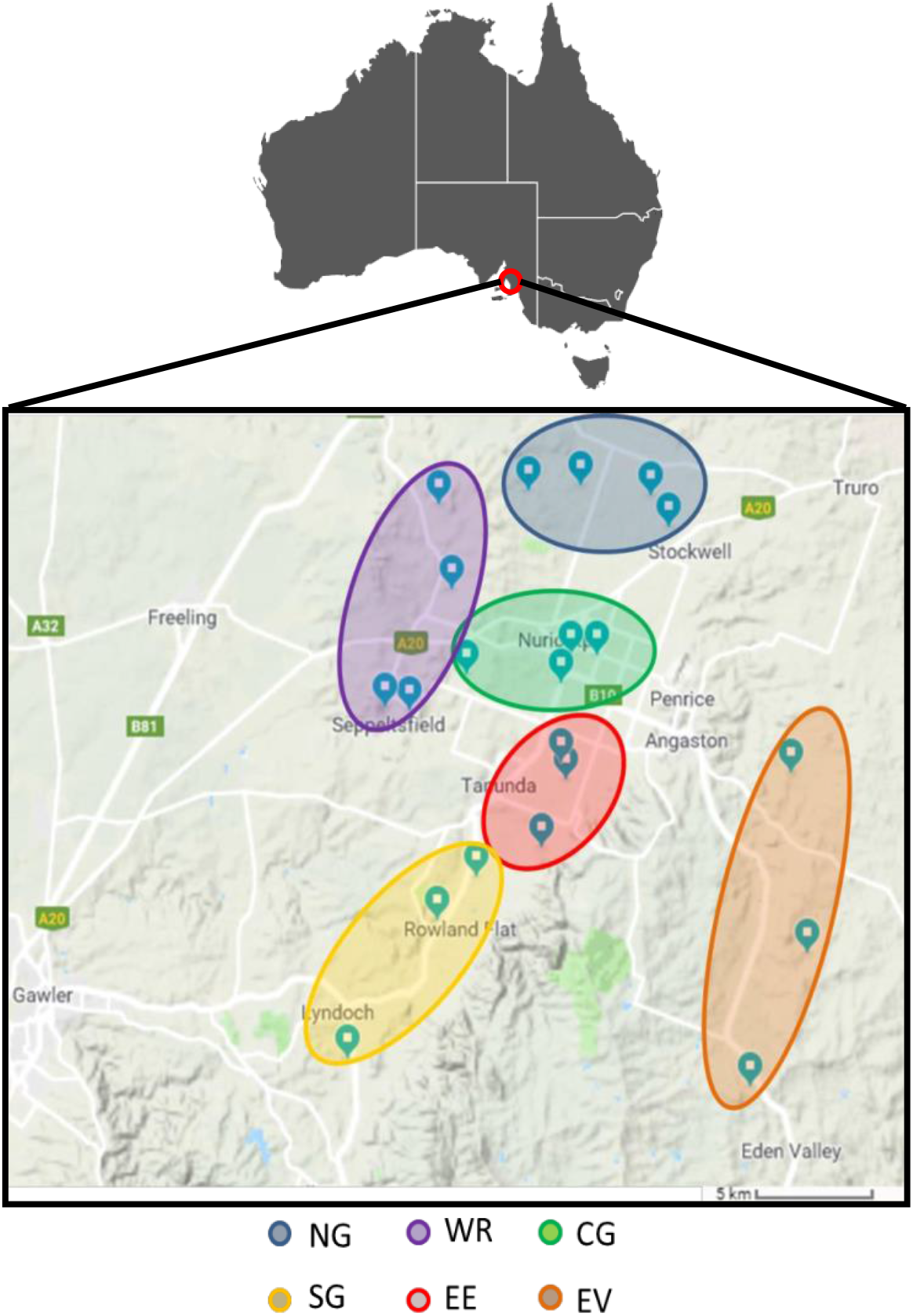
Geographical location of grapevine tissue samples analysed in this study. Leaf and berry samples were harvested from selected vineyards from the Barossa wine region, Australia^34^. Northern Grounds (NG, blue) Central Grounds (CG, green), Southern Grounds (SG, yellow), Western Ridge (WR, purple), Eastern Edge (EE, red) and Even Valley (EV, orange).

### Identification of putative dicistronic tRNA-mRNA transcripts

We searched for combinations of tRNA and adjacently located protein coding mRNA genes that were expressed forming one continuous transcript. With that objective, we developed DiRT, a bioinformatic pipeline to systematically analyse high-throughput, short read-based RNA-sequencing data for actively transcribed tRNA-mRNA dicistronic loci (Figure 2). The pipeline identifies dicistronic transcripts in both mRNA-tRNA and tRNA-mRNA genomic orientation, however for the sake of simplicity, both dicistronic combinations will be hereafter be referred to as tRNA-mRNA transcripts. The pipeline takes into consideration reads mapping in the tRNA, mRNA and the intervening intergenic region to predict dicistronic tRNA-mRNA candidates. Biological replicates were used to estimate background noise and improve the accuracy of the predictions.

**Figure 2.**
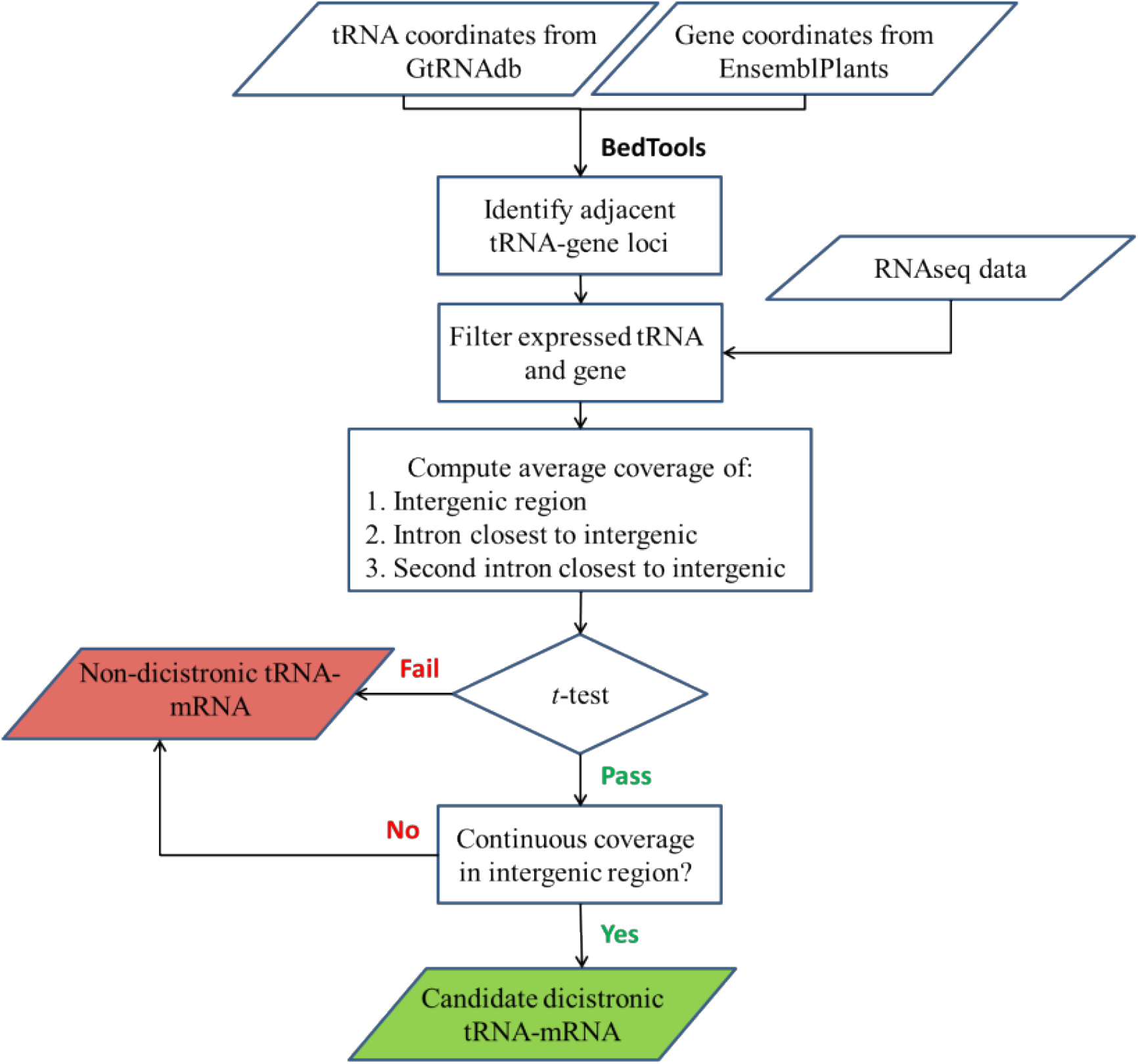
Schematic workflow of the DiRT pipeline to identify dicistronic candidates from RNA-seq data. tRNA gene and protein coding gene coordinates were retrieved from GtRNAdb and Ensembl Plants respectively. tRNA-protein coding gene pairs that occupied contiguous spaces in the genome were assembled and selected for subsequent analysis if they were transcribed based on RNA-seq reads (Raw read >= 1 for tRNAs and raw read >= 10 in PCGs). Dicistronic tRNA-mRNA are identified by assessing active transcription of the intergenic region. Candidate tRNA-mRNA are selected if the intergenic regions had significantly higher expression (FDR < 0.05) than the closest two introns. Pairs of tRNA-gene are classified as putatively dicistronic tRNA-mRNA if the intergenic region showed continuous sequencing coverage between the tRNA and the mRNA of the protein coding gene.

The Genomic tRNA Database predicts 609 tRNA genes in the *V. vinifera* genome based on the tRNAscan-SE tool^19^. From these, 116 tRNA genes overlapped with protein coding genes (PCGs), defined by genomic boundaries that extend between the 5’ and 3’ untranslated regions. tRNA genes that overlap with the PCGs (5’ or 3’ untranslated regions or introns) were removed from further analysis since such reads could not be unambiguously assigned to either the tRNA or the PCG. Using DiRT, we detected 137 transcribed tRNA genes (read count ≥ 1) in both tissue samples (124 in leaf and 90 in berry) across all sub-regions. From the 137 tRNA genes transcribed, 77 were expressed in both tissues, 47 tRNAs were leaf specific and 13 tRNAs were berry specific. From the expressed tRNAs, 136 presented typical tRNA secondary structure with associated anticodon and just one tRNA from leaf was classified as undefined (*Vitis vinifera* 12x reference genome, PN40024). In general, the numbers of expressed tRNAs are underrepresented in a standard short-read RNA-seq library due to tRNA secondary structure and post-transcriptional modifications limiting detection^20^. Although underrepresented, we evaluated the sensitivity of the pipeline to identify dicistronic tRNA-mRNA transcripts using a standard RNA-seq workflow as it permits expression analysis of monocistronic and dicistronic transcripts from the same sequenced library. Individual tRNA genes detected displayed a wide-range of transcript abundances covering the 20 isoacceptor families in both leaves and berries, showing a distinct tRNA expression profile across the six regions analysed (Supplemental Fig S1 and Supplemental Table S2). We assembled combinations of tRNA-PCGs and identified 81 expressed tRNA-mRNA combinations (Figure 3a) in leaves and 50 in berries. As the intergenic region between the transcribed tRNA and mRNA for sequence reads would be indicative of co-transcription, we tested the significance of reads in the intergenic region to eliminate background noise attributable to DNA contamination or spurious transcription events that would not be observed in biological replicates. tRNA-mRNA combinations were selected for further analysis only if the coverage of their intergenic region was significantly higher (t-test, FDR < 0.05) than reads detected in the two closest introns (Figure 3b and 3c). Finally, candidates that passed both tests were tested for continuous read coverage in the intergenic region indicating transcriptional read-through of the region between the tRNA and the mRNA (Figure 3d). DiRT identified 16 dicistronic tRNA-mRNA transcripts in leaves and nine in berries, of which six were present in both tissues (Table 1) across 13 of the 19 *V. vinifera* chromosomes. Sequencing coverage was significantly higher (t-test, FDR < 0.05) in intergenic regions than in the first two introns of dicistronic tRNA-mRNA pairs. Conversely, no significant difference in coverage was observed for tRNA-mRNA pairs deemed non-dicistronic and these transcripts had expected read coverage breakpoints in the intergenic region between tRNA and mRNA (Supplemental Fig S2).

**Figure 3:**
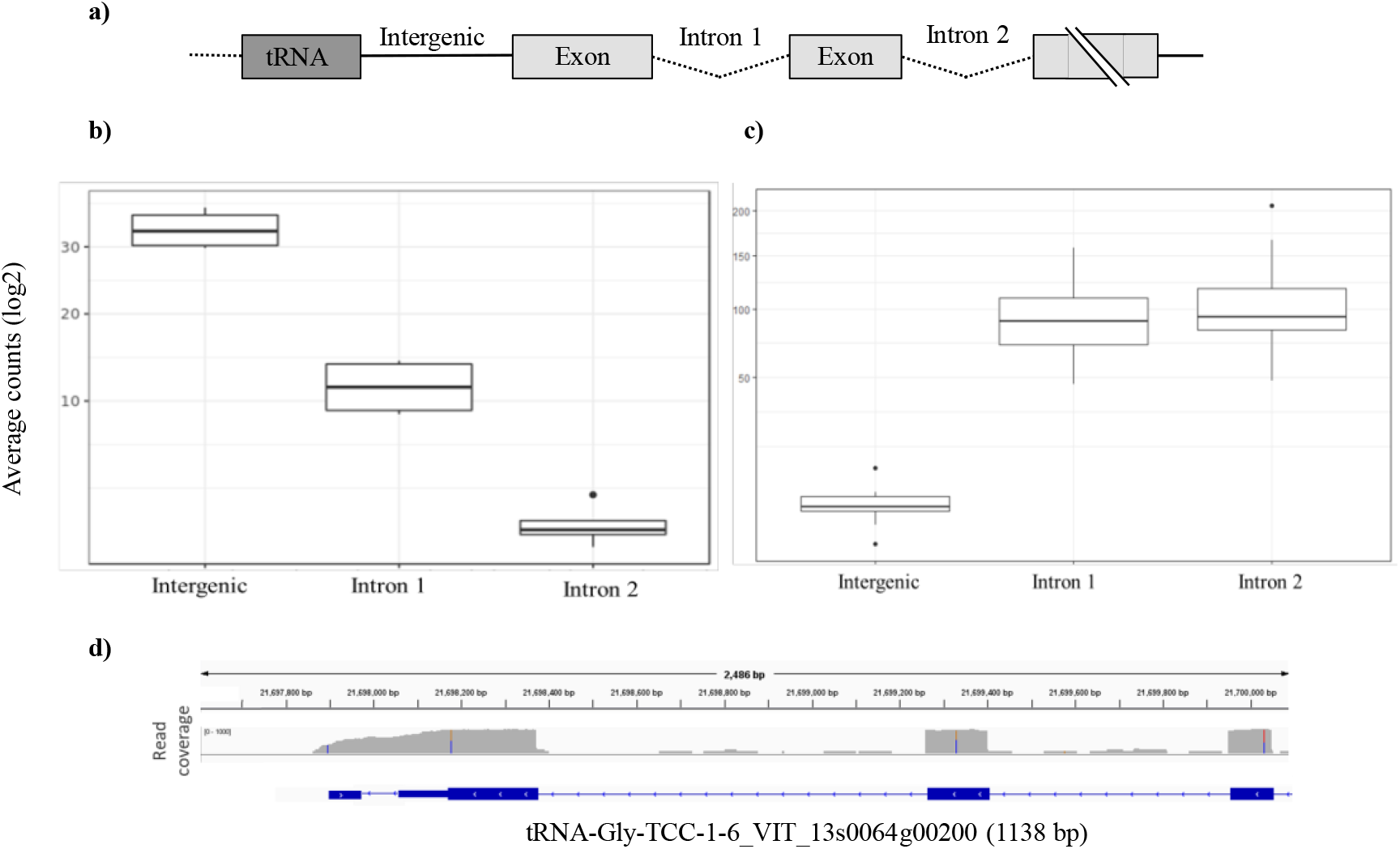
Assessing active transcription of intergenic region of putative dicistronic tRNA-mRNAs transcripts from RNA-seq data. a) Gene model of dicistronic tRNA-mRNA transcript in which the intergenic region was expressed b) Average read coverage of the intergenic region versus the closest two introns of a tRNA-mRNA combination that passed b) (tRNA^Gly-GCC-1-6^ VIT_08s0058g00460) and failed c) (tRNA^Tyr-GTA-4-1^ VIT_00s0505g00030) the t-test (p-value < 0.05). d) Genome browser view of a candidate dicistronic tRNA-mRNA formed by tRNA^Gly-TCC-^ 1-6 and VIT_13s0064g00200 identified using the DiRT pipeline.

**Table 1.**
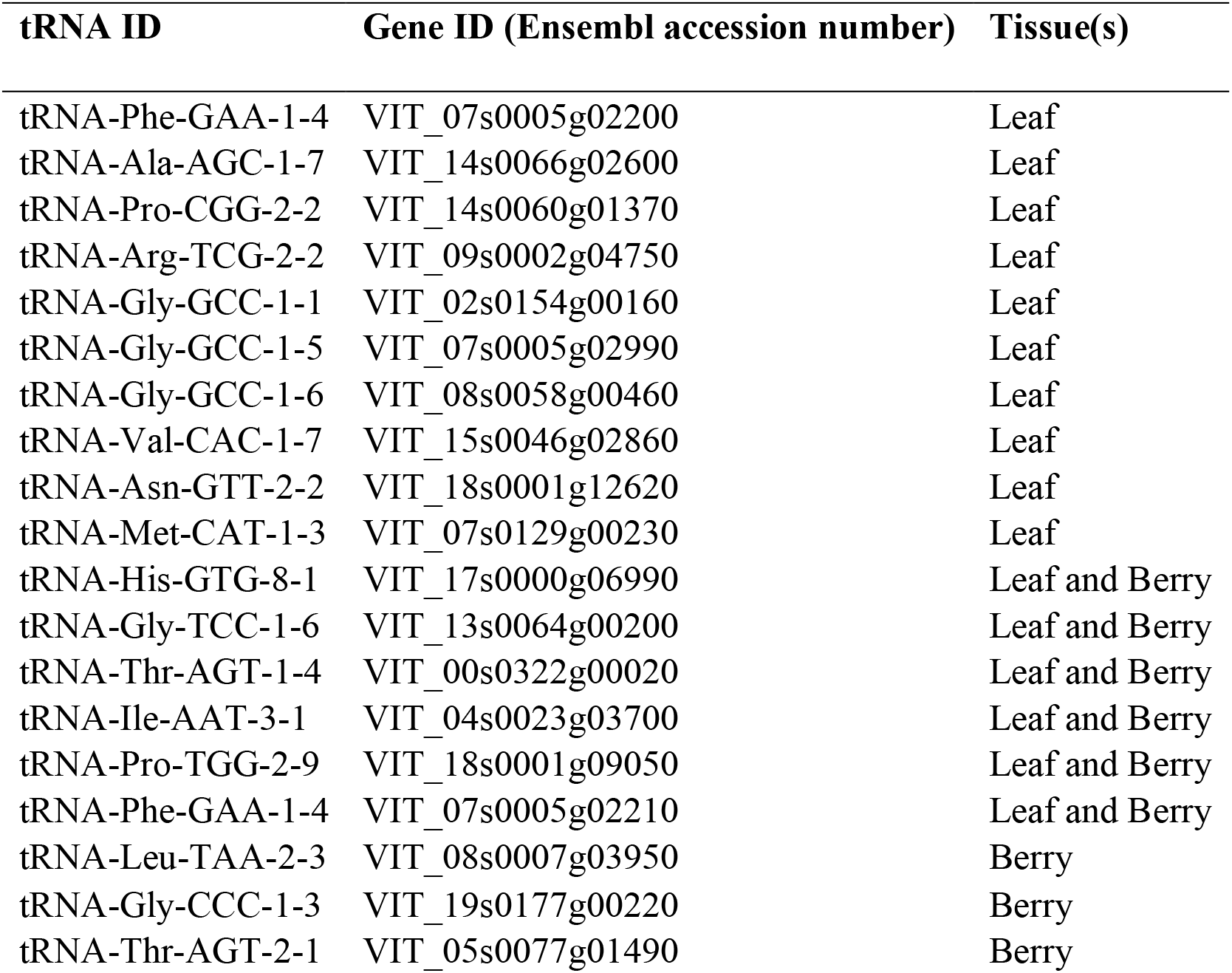
Dicistronic tRNA-mRNA candidates identified from RNA-seq data in leaves and berries of grapevine.

| tRNA ID | Gene ID (Ensembl accession number) | Tissue(s) |
| --- | --- | --- |
| tRNA-Phe-GAA-1-4 | VIT_07s0005g02200 | Leaf |
| tRNA-Ala-AGC-1-7 | VIT_14s0066g02600 | Leaf |
| tRNA-Pro-CGG-2-2 | VIT_14s0060g01370 | Leaf |
| tRNA-Arg-TCG-2-2 | VIT_09s0002g04750 | Leaf |
| tRNA-Gly-GCC-1-1 | VIT_02s0154g00160 | Leaf |
| tRNA-Gly-GCC-1-5 | VIT_07s0005g02990 | Leaf |
| tRNA-Gly-GCC-1-6 | VIT_08s0058g00460 | Leaf |
| tRNA-Val-CAC-1-7 | VIT_15s0046g02860 | Leaf |
| tRNA-Asn-GTT-2-2 | VIT_18s0001g12620 | Leaf |
| tRNA-Met-CAT-1-3 | VIT_07s0129g00230 | Leaf |
| tRNA-His-GTG-8-1 | VIT_17s0000g06990 | Leaf and Berry |
| tRNA-Gly-TCC-1-6 | VIT_13s0064g00200 | Leaf and Berry |
| tRNA-Thr-AGT-1-4 | VIT_00s0322g00020 | Leaf and Berry |
| tRNA-Ile-AAT-3-1 | VIT_04s0023g03700 | Leaf and Berry |
| tRNA-Pro-TGG-2-9 | VIT_18s0001g09050 | Leaf and Berry |
| tRNA-Phe-GAA-1-4 | VIT_07s0005g02210 | Leaf and Berry |
| tRNA-Leu-TAA-2-3 | VIT_08s0007g03950 | Berry |
| tRNA-Gly-CCC-1-3 | VIT_19s0177g00220 | Berry |
| tRNA-Thr-AGT-2-1 | VIT_05s0077g01490 | Berry |

In total, 19 individual tRNA genes, representing 13 isoacceptor families were found to be dicistronic with the neighbouring protein coding genes, among which, glycine tRNA genes were the most common. We validated the expression of the tRNA-mRNA dicistronic candidates identified by DiRT through RT-PCR. We tested candidates with tissue-specific expression (leaf: tRNA^ValCAC^-VIT_15s0046g02860 (Figure 4a), berry: tRNA^GlyCCC1.3^-VIT_19s0177g00220 (Supplemental Figure S3a)) and a candidate that was expressed in both tissues (tRNA^ProTGG2.9^-VIT_18s0001g09050 (Figure 4b and Supplemental Figure S3b).

**Figure 4.**
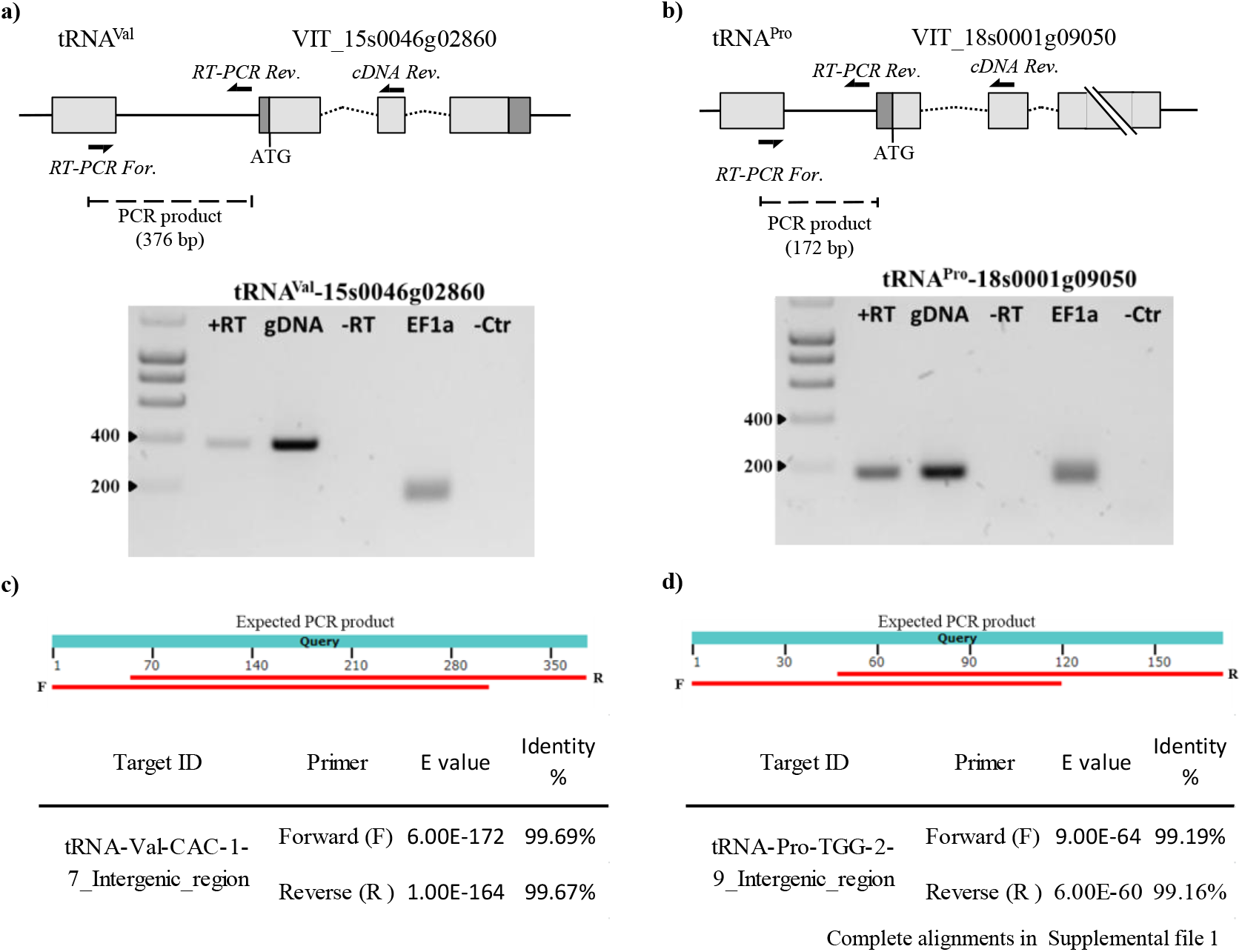
RT-PCR confirmation of identified dicistronic transcripts in leaf samples. Model of putative dicistronic tRNA-mRNA transcript showing primers used for cDNA synthesis (cDNA rev.) and for the PCR reaction (RT-PCR For and RT-PCR Rev). Confirmation of actively transcribed intergenic region through RT-PCR for candidates a) tRNA ^ValCAC^-VIT_15s0046g02860 (376 bp) and b) tRNA^ProTGG^-VIT_18s0001g09050 (172 bp). +RT: cDNA as template, gDNA: genomic DNA was used as a control, -RT: RT-PCR negative control, EF1a: Elongation Factor 1-alpha was used as a positive control (150 bp), -Ctr: PCR negative control. Alignment of the sequenced PCR product for candidate c) tRNA^ValCAC^-VIT_15s0046g02860 and d) tRNA^ProTGG^-VIT_18s0001g09050 to the expected PCR product confirmed active transcription of the intergenic region.

To confirm the dicistronic nature of the transcript, the intergenic region spanning between the tRNA and protein coding gene was PCR amplified using leaf and berry cDNA as template. For the candidates tested, a single band of the expected product size was obtained. Sanger sequencing of the PCR product confirmed the amplification of the intergenic regions (Figure 4c, Figure 4d and Supplemental Table S3).

### Characteristics of grapevine dicistronic tRNA-mRNA candidates

The genomic distance between expressed tRNA and PCGs that formed dicistronic transcripts was no longer than 1065 base pairs (bp), with a median intergenic distance of 133 bp (Supplemental Fig S4). The observed frequency of mRNAs forming dicistronic transcripts decreased with distance both upstream and downstream from the tRNA component of the dicistronic pair. We next analysed the upstream and downstream sequences of the dicistronic tRNA in search of *cis*-acting signals that might explain transcriptional read-through to the adjacent PCG. Sequence analysis of 20 bp upstream and downstream of the dicistronic tRNA revealed the presence of canonical motifs associated with tRNA transcription efficiency (Supplemental Fig S5)^6,21^. This included a high proportion of A nucleotides upstream of the transcription start site, important for maintaining high tRNA expression, and a short stretch of downstream T nucleotides for RNA Polymerase III transcription termination. We did not identify any novel conserved sequence between the dicistronic candidates that could act as a mediating signal for the co-transcription of the tRNA and PCG.

When we compared the expression of both mRNAs and tRNA deemed to be dicistronic in this study against the background of all expressed genes, we found that dicistronic tRNA-mRNAs’ expression did not correlate with high abundance genes in either leaf or berry tissue (Supplemental Fig S6). Most values of gene and tRNA expression were between the 25^th^ and 75^th^ % of the distribution of the total gene expression.

Of the nineteen PCGs that formed dicistronic transcripts, fourteen have annotated functions and five are described as uncharacterised in the EnsemblPlants release 45 database^22^ (Supplemental Table S4). Six of the fourteen characterised genes are associated with functions relating to nucleic acid binding or processing activity and three are involved in the flavin biosynthesis pathway. A BLAST search in the *Arabidopsis thaliana* genome revealed 11 of the 19 *Vitis* dicistronic PCGs have a closely related *A. thaliana* ortholog that is either dicistronic (4/11)^7^ or the mRNA has been demonstrated to be mobile (8/11, PlaMoM database)^23^ (Supplemental Table S4). Notably, the common *A. thaliana* dicistronic PCGs are also co-transcribed with the same tRNA isodecoder as in grapevine.

### Regional patterns of dicistronic expression

We next assessed if the geographical origin of the samples had an effect on the expression of dicistronic transcripts. We first analysed the expression of all tRNAs expressed in each tissue and we used hierarchical clustering to group sub-regions according to their tRNA expression patterns in leaves and berries. Both tissues presented two main clusters containing three sub-regions each (Figure 5). The tRNA expression in Eastern Edge and Northern Grounds clustered together in both tissues, while the clustering of the four other subregions were tissue dependent. We then analysed the expression of the tRNA genes, the intergenic regions and PCGs forming dicistronic transcripts independently. We used RNA-seq reads mapping specifically to the intergenic region as a proxy to estimate expression patterns of dicistronic candidates, as reads mapping to the flanking tRNA and PCG loci could originate from both monocistronic and dicistronic transcripts (Figure 5). Sub-regional clusters for tRNAs forming part of dicistronic constructs were similar to those observed for all expressed tRNAs in both tissues (Figure 5). In leaf, one of the main clusters (SG, EE and NG) was the same for all expressed tRNAs and tRNAs that were part of dicistronic constructs. While in berry, EE/NG and CG/WR clusters were the same in all expressed tRNAs and dicistronic tRNAs. EE/NG and CG/WR clustered together in both tissues, while SG and EV clustering was tissue dependant. When the expression of the intergenic regions and dicistronic PCGs was used rather than tRNA, sub-regional clustering was tissue and dicistronic construct component specific (intergenic region or PCG).

**Figure 5.**
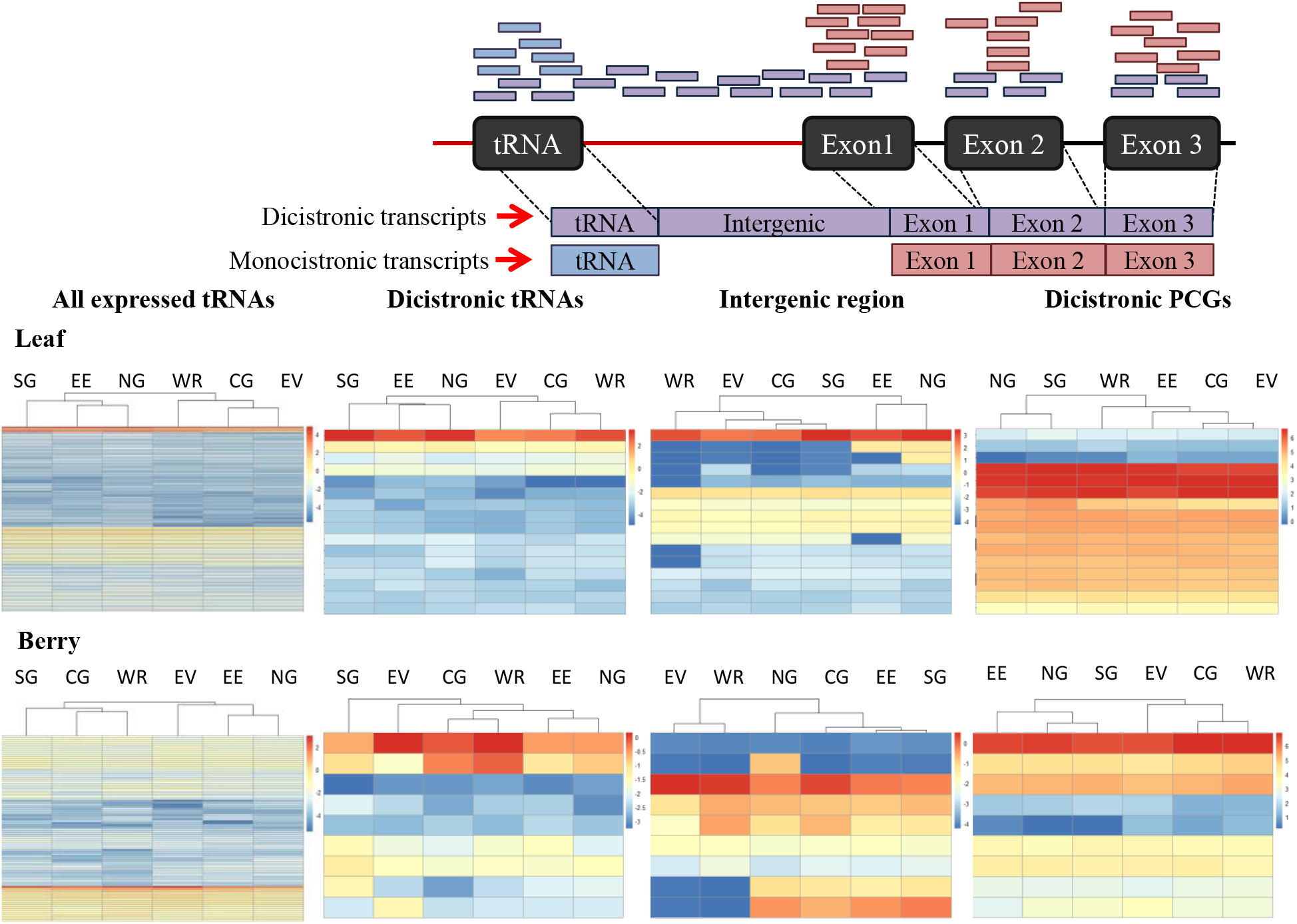
Effect of region of origin on the expression of dicistronic tRNA-mRNAs. Top panel: Schematic representation of RNA-seq reads mapping when originated from a dicistronic transcript (purple bars), a monocistronic tRNA transcript (blue bars) and a monocistronic protein coding gene (red bars). Bottom panel: Heatmap of the expression (logCPM) of all expressed tRNAs, dicistronic tRNAs, intergenic region and dicistronic protein coding genes (rows) for each sub-region from the Barossa Wine growing region (columns) for leaf and berry samples. Dendrograms represent the hierarchical clustering analysis of the sub-regions according to each genomic feature expression pattern.

We analysed the expression patterns of all tRNA expressed and the dicistronic construct components (i.e. dicistronic tRNAs, PCG and intergenic regions) through local Fisher discriminant analysis (LFDA). We observed that the expression patterns of PCGs occupies a unique eigen space, while dicistronic tRNAs and intergenic regions shared the eigen space occupied by all expressed tRNAs (Supplemental Fig S7). Consistent with this observation, correlation analyses of the expression of the different part of the candidates dicistronic transcripts showed that the absolute values of Pearson correlation coefficients were generally higher between the expression of dicistronic tRNAs and the expression of the intergenic region than between the expression of PCGs and the expression of the intergenic regions on both tissues (Supplemental Table S5). These correlations were only significant (Pearson correlation, p-value < 0.05) between dicistronic tRNAs and intergenic regions in leaves (Supplemental Table S5).

### Evolutionary co-appearance of dicistronic tRNA-mRNA transcripts and vasculature in land plants

The expression of the tRNA-mRNA dicistronic transcripts in grapevine leaf and berry tissues and across multiple growing regions indicates tissue-specific function and environmental regulation. In addition, the earlier report in *Arabidopsis* and tobacco of dicistronic tRNA-mRNA transport through phloem suggests these transcripts are enriched in vascular tissue potentially serving as long-distance signalling molecules^7^. To gain further insights into the signalling potential of these transcripts across the plant kingdom, we next investigated if the emergence of tRNA-mRNAs as dicistronic transcripts co-occurred with the appearance of vasculature during the evolution of land plants. To test this hypothesis, the DiRT pipeline was used to identify dicistronic tRNA-mRNAs from RNA-seq datasets across a range of flowering plants, (*Arabidopsis thaliana, Vitis vinifera, Oryza sativa* and *Brachypodium distachyon*), ancient vascular plants (*Azolla filiculoides* and *Salvinia cucullata* (ferns) and *Selaginella moellendorffii* (lycophytes)), and non-vascular species (*Physcomitrella patens* (mosses) and *Marchantia polymorpha* (liverworts)). RNA-seq data for each representative vascular and non-vascular species was accessed from the short-read archive (SRA) and filtered based on organ samples that included vasculature tissue and sequencing criteria to ensure consistency across the libraries analysed (See Methods section). In total, 149 Illumina-based RNA sequencing libraries from the nine vascular and non-vascular species were analysed using the DiRT pipeline (Table 2).

**Table 2.** Multi-species detection of dicistronic tRNA-mRNA transcripts using DiRT to analyse publicly available RNA-seq experiments accessible from the Sequence Read Archive (SRA).

| Species | SRA Project number | RNA-seq libraries analysed (average reads/library) | Tissue | Number of tRNA isodecoders forming dicistronic transcripts | Number of dicistronic tRNA-mRNA candidates identified |
| --- | --- | --- | --- | --- | --- |
| *Oryza sativa* ssp. *japonica* | PRJNA403975<br>PRJNA301554<br>PRJNA350792 | 17 (67 M) | leaf,<br>root | 14 | 29 |
| *Brachypodium distachyon* | PRJNA293888<br>PRJNA371381<br>PRJNA383406 | 12 (57 M) | leaf,<br>root | 2 | 3 |
| *Arabidopsis thaliana* | PRJNA271927<br>PRJNA241198<br>PRJNA246731<br>PRJNA318918 | 24 (46 M) | leaf,<br>root,<br>shoot | 9 | 11 |
| *Vitis vinifera* | PRJNA311966<br>PRJNA383160<br>PRJNA342391<br>PRJNA260535<br>PRJNA307079<br>PRJNA386484<br>PRJNA433817<br>PRJNA268857 | 44 (30 M) | leaf,<br>root,<br>berry | 15 | 39 |
| *Azolla Filiculoides* | PRJNA430527<br>PRJNA264391 | 9 (40 M) | whole plant,<br>root | 2 | 4 |
| *Salvinia cucullata* | PRJNA430459 | 6 (43 M) | leaf | 1 | 2 |
| *Selaginella moellendorffii* | PRJEB29012<br>PRJNA271595 | 9 (31 M) | leaf,<br>root | 23 | 49 |
| *Physcomitrella patens* | PRJNA417704<br>PRJNA397394<br>PRJNA294412<br>PRJNA259146<br>PRJNA259147 | 17 (32 M) | whole plant,<br>protonema,<br>chloronemata | 0 | 0 |
| *Marchantia polymorpha* | PRJDB4420<br>PRJNA433456 | 11 (21 M) | thallus,<br>antheridiophore,<br>archegoniophore | 0 | 0 |

Dicistronic tRNA-mRNA transcripts were identified in all taxa presenting vascularization. In total, 139 dicistronic tRNA-mRNA transcripts were detected across eudicots, monocots, fern species and the lycophyte representative, *Selaginella moellendorffii* (Table 2). Corroborating evidence was also obtained in *Vitis vinifera*, with over half of the tRNA-mRNA candidates identified from Barossa valley leaf and berry tissue sampled as part of this study were also identified as dicistronic from the publicly available *Vitis* RNA-seq datasets (10/19 candidates). Across all vascular species analysed, 66 tRNA genes were detected to be co-transcribed with the neighbouring protein-coding gene, representing 20 distinct isoacceptor families (41 anticodons) (Figure 6a). We did not observe overrepresentation of any one tRNA isoaceptor family that forms dicistronic transcripts across all species, however, tRNA^Glu(CTG)^ was observed to be co-transcribed with a PCG in all four flowering plants analysed.

**Figure 6.**
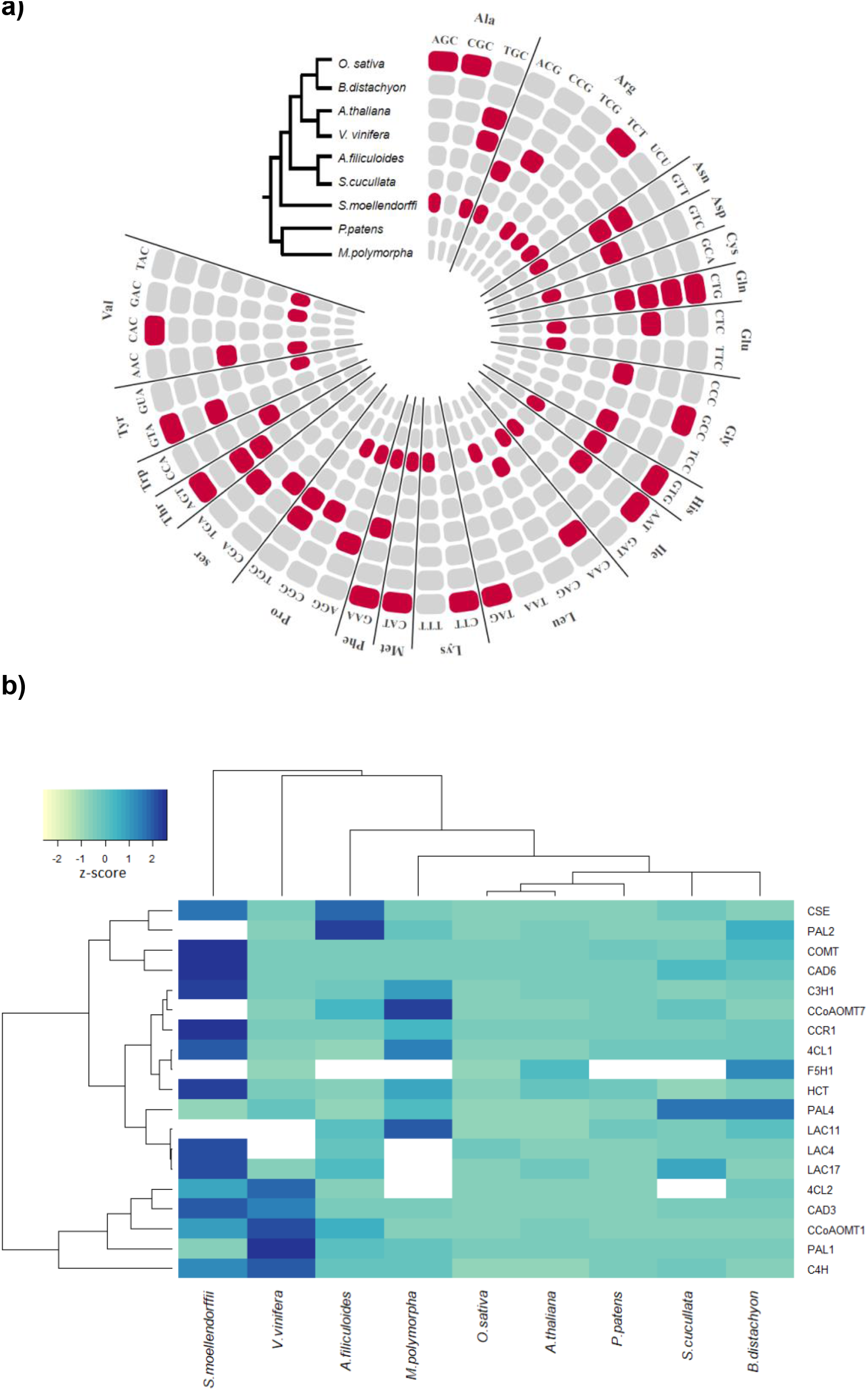
Identification of dicistronic tRNAs in land plants using the DiRT pipeline. a) circular visualisation of plant tRNAs co-transcribed with PCGs in land plants using publicly available RNA-seq datasets accessible from the SRA. Concentric circles represent tRNAs detected as dicistronic (red) or non-dicistronic (grey) in one or more RNA-seq datasets for each plant species analysed. In each circle, individual tRNAs are organised alphabetically by amino acid and are aligned for each species based on the anticodon. Details of the number of times a specific tRNA was detected as dicistronic in RNA-seq experiments and the co-transcribed PCG are provided in Supplemental Table S11. b) RNA-seq expression heat map (TPM) of orthologous lignin biosynthesis genes involved in vasculature development. Average TPM expression values were derived from the same RNA-seq datasets used for analysing dicistronic tRNA-mRNA transcription. Colour represents the expression Z-score (TPM minus mean over s.d) of each gene (row) in a given species (column). A white panel is used to indicate cases where unequivocal ortholog assignment was not possible or where an orthologous gene member could not be identified.

Dicistronic transcripts were reproducibly detected in multiple and independent RNA-seq experiments from vascular species. In contrast, we found no evidence of these transcripts in early lineages of extant land plants *Physcomitrella patens* and *Marchantia polymorpha* that lack vascular tissue (Figure 6a). Using the same RNA-seq datasets, we also analysed expression of a core set of vasculature genes involved in lignin biosynthesis and secondary wall biosynthesis to ensure vascular tissues were being represented in all RNA-seq library samples chosen in the analysis. Expression of many, but not all, orthologous genes involved in lignin and secondary cell wall biosynthesis was detected in both vascular and non-vascular plants confirming earlier observations that much of the genetic pathway for vasculature development was established early in lower land plants and developed during the course of evolution^24,25^ (Figure 6b, Supplemental Fig S8).

## Discussion

In this study, using an RNA-seq approach, we found that 13.9% (19/137) of all expressed tRNAs in grapevine leaf and berry samples were putatively expressed in a dicistronic manner, with neighbouring protein coding genes. We developed DiRT, a customised, computational pipeline to specifically detect dicistronic tRNA and mRNA candidates using stringent criteria. Using DiRT we were able to identify dicistronic transcripts in two different grapevine tissues (i.e. leaf and berry) sampled from commercial vineyards. Validation of the pipeline to accurately predict dicistronic candidates was confirmed in both tissues through RT-PCR detection and Sanger sequencing of dicistronic candidates in leaf samples.

Interestingly, of the 12 tRNA isoacceptor families (representing 15 distinct anticodons) found to be dicistronic in *Vitis vinifera*, 11 tRNA families have also found to be dicistronic in *A. thaliana*, and associated with transcripts demonstrated to be mobile between roots and shoots^7,26^. Among these tRNA coding for Gly^GCC^ and Met^CAT^ were able to mobilise mRNA transcripts to different tissues as part of a fusion construct and translate into functional proteins in grafted *A. thaliana* plants, indicating that these tRNA were able to confer mobility to these transcripts^7^. This suggests a non-autonomous role for dicistronic tRNAs in delivering mRNA transcripts to distantly located tissues. A recent study also revealed that mobile RNA transcripts are enriched in the modified base 5-methylcytosine (m^5^C), indicating a role of RNA cytosine methylation in systemic RNA movement^27^. In plants, tRNA and mRNA m^5^C methylation is mediated by the methyltransferases DNMT2 and TRM4B^20,28,29^ and loss of these enzymes was demonstrated to impair transcript mobility^27^. Future studies will need to be undertaken to determine if the dicistronic tRNAs identified in this study also confer mRNA mobility and to assess the role of cytosine methylation in mRNA transport in grapevine.

For four of the 19 dicistronic candidates we also observed sequence conservation between *A. thaliana* and *V. vinifera* for the protein coding gene and the adjacently co-transcribed tRNA genes. This indicates the genomic clustering of tRNA and protein coding gene is, in some cases, evolutionarily conserved between both species. The dicistronic activity at these conserved loci may provide an explanation of why such syntenic clusters are conserved through evolution and suggests that these transcripts may have an important functional role.

Of the 19 dicistronic tRNA genes identified in the *Vitis vinifera* genome, 18 were located fewer than 1000 base pairs from the co-transcribed upstream or downstream protein coding gene (median distance 133 bp). Our findings suggest that tRNA genes and protein coding genes need to be closely positioned in the genome in order to form dicistronic transcripts. Similar observations were obtained in *A. thaliana*, where the majority of the previously identified PCGs forming part of mobile dicistronic transcripts were located less than 200 bp from their partner tRNA^7^. Genomic proximity was found to be even closer between co-transcribed tRNA and snoRNA genes identified in higher plants such as *A. thaliana, M. truncatula, P. trichocarpa, O. sativa* and *B. distachyon*, in which the intergenic region ranged between 1 to 16 base pairs^6^.

Previous studies have indicated that a large proportion of mobile transcripts are also highly abundant^26,30^. This suggests that passive diffusion of these transcripts through the phloem may contribute to their mobility. A significant proportion (11.4%) of these transcripts was subsequently shown to be dicistronically associated with tRNA^7^. However, when we assessed the expression levels of mRNA and tRNA that formed dicistronic transcripts in grapevine, we did not observe higher abundance of these transcripts in either tissue analysed. Thus, in our study, the expression level of the tRNA and mRNA was not a good indicator of the formation of dicistronic transcripts. In eukaryotes, tRNA and mRNA are transcribed by different types of RNA polymerase. RNA polymerase II (Pol II) transcribes protein coding genes and RNA polymerase III (Pol III) for a variety of genes that generally encode for RNAs with catalytic activity such as tRNA^31^. Results from Kruszka et al. ^5^ suggested that, in *A. thaliana*, dicistronic tRNA-snoRNA are transcribed by Pol III from the tRNA gene promoter. However, Pol III transcribes genes shorter than 400 base pairs^31^ and the dicistronic transcripts identified in our study were considerably longer (between 1486 to 6002 bp) suggesting Pol III may not be co-transcribing these transcripts. A comparative analysis of flowering species showed a poly-T stretch immediately downstream of ≥ 90% of tRNA genes^6^. Additionally, Michaud et al.^6^ reported that the few tRNAs lacking poly-Ts were capable of forming dicistronic transcript with snoRNAs. The authors hypothesized that the lack of the poly-T transcriptional termination signal could be a possible explanation for why these transcripts were transcribed as a single unit by Pol III. Sequence analysis of the upstream and downstream sequences of the dicistronic tRNAs identified in our study revealed canonical elements previously associated with transcription start and termination^6,21^. In particular, all dicistronic tRNA transcripts we identified had a poly-T termination signal suggesting that the transcriptional mechanism of these transcripts may be different from tRNA-snoRNAs. Another possibility is that the tRNA and mRNA are transcribed independently as monocistronic transcripts and are then ligated or spliced together to form a dicistronic transcript. Currently, the DiRT pipeline cannot distinguish between transcriptional read-though and post-transcriptional ligation of two independent transcripts. However, as both possibilities result in the same effective dicistronic molecule, detection is nevertheless important for understanding the physiological functions of these transcripts. The expression patterns of all genomic features studied (i.e. tRNAs, PCGs, and intergenic regions (considered a proxy for dicistronic tRNA-mRNA transcripts)) were found to be organ specific and sensitive to regional environmental differences. The effect of organ and environment on PCG and tRNA gene expression has been extensively studied in grapevine^9,15^. Special effort has been put into deciphering the effect that the growing environment has on fruit quality traits associated with wine regionality^32,33^. However, the effect that the environment and tissue have on tRNA expression and on dicistronic transcript expression has not been previously described. Our results show that the expression patterns of dicistronic transcript-forming tRNA genes mimic those of all expressed tRNA (Supplemental Fig S7). We also found that the expression of dicistronic tRNA-mRNAs, measured as the expression of the intergenic region, showed a higher correlation with that of dicistronic tRNA than with that of dicistronic PCGs in both tissues. Although this correlation was only statistically significant in leaves, the lack of statistical significance in berry samples could be due to the low number of dicistronic transcripts identified in berries compared to leaves (9 vs 16 respectively). However, berries in bunches undergoing veraison present high levels of phenological variability. Such variability, if not properly captured, can influence significantly the outcome of transcriptomic studies, as shown previously^34–36^. Taking this in consideration, and although our sampling strategy was designed to capture vineyard variation, we cannot rule out that the differences in dicistronic transcript expression observed between regions are not due to the intrinsic variability imposed by veraison.

Taken collectively, our results suggest that environmentally induced dicistronic tRNA-mRNA expression is, at least partially, directed by the mechanisms regulating tRNA expression. Prior work in the model species *Arabidopsis thaliana* and tobacco, demonstrate tRNAs and closely related tRNA-like structures that are co-transcribed with mRNA have the ability to be systemically transported via the phloem vasculature. The translation of the mobile mRNA product into a functional protein indicates that such molecules have the potential to act as non-autonomous long-distance signals^7^. To provide a broader context for these transcripts in plants, we show here the first evidence for dicistronic tRNA-mRNA transcription in a commercially important crop species grown in field conditions. Furthermore, through a multi-species RNA-seq transcriptome analyses, we demonstrate wide prevalence of dicistronic tRNA-mRNA in land plants. Importantly, the detection of these transcripts in flowering angiosperms, ferns and lycophytes but absence in basal plant lineages such as liverworts and mosses indicates that the emergence of these co-transcribed molecules coincided with evolution of the vasculature trait in land plants. As the vascular system evolved in plants as means to translocate water, nutrients and other photoassimilates facilitating terrestrial colonization, the concurrent appearance of dicistronic tRNA-mRNA transcripts reinforces their potential role in long-distance communication. Thus, the evolution of vasculature may have led to plants acquiring a new role for tRNAs in mediating transport of large mRNA transcripts to distant organs as a means to integrate developmental and environmental cues.

We detected a varying number of dicistronic tRNA-mRNA transcripts in the seven representative vascular species analysed ranging from 2 in *S. cucullata* to 49 in *S. moellendorfii*. Such variation could reflect species specific differences of tRNA-mediated mRNA transport though the vasculature, varying number of tRNA genes in the genome and/or differences in co-transcriptional machinery. Given the paucity of knowledge regarding tRNA-mRNA transcripts, biological differences between species are difficult to determine at this stage. Moreover, although we controlled for the expression of vascular genes and mined the range of high-quality datasets available publicly, we cannot rule out that subtle differences in tissue type and sampling conditions in each species analysed may have contributed to the variation of discistronic transcripts observed. For the purposes of this study however, the results from the multi-species comparison served as qualitative measure of dicistronic tRNA-mRNA transcription in vascular versus non-vascular species rather than as quantitative study. Future work comparing syntenic regions of tRNA and mRNA genes between species and analysing the biological relevance of transported mRNAs will help in understanding species-specific functions of dicistronic transcripts.

It remains to be elucidated if tRNA-mRNA movement requires RNA-binding proteins for vascular transport, as is the case for viruses and viroids that require movement proteins for RNA transport. Future studies that use our *in silico* approach to survey expression patterns of these transcripts in an even wider range of model and non-model plant species and across a range of tissues will shed more light on expression dynamics; the source, intermediate and target cell types and signalling function of these transcripts.

## Material and methods

### Sampling material

Tissues were sampled from own-rooted grapevines (*Vitis vinifera*) cv. Shiraz from 22 commercial vineyards located in the Barossa wine zone (South Australia, Australia). Vineyards were selected as part of a larger study of Barossa Terroir^37^. Vineyards were chosen to be representative of the climate, soil and management practices that are used in the different Barossa sub-regions. These sub-regions are the Eden Valley (EV) (3 vineyards), Northern Grounds (NG) (4 vineyards), Central Grounds (CG) (4 vineyards), Southern Grounds (SG) (3 vineyards), Eastern Edge (EE) (4 vineyards) and Western Ridge (WR) (4 vineyards).

Leaves and berries were collected from nine plants in each of three rows in each vineyard (total of 198 plants) during the 2016 growing season. The first fully expanded leaf at budburst (E-L 7) was collected from three nodes per plant. Nine berries were collected at veraison (E-L 35) from three different bunches per plant (i.e. three berries per bunch). All samples were taken before dawn (between 10:00 pm and sunrise) to minimise variability associated with differences in plant water status^38^. Samples were snap-frozen in liquid nitrogen in the vineyard of collection. All samples of the same organ (leaf or berry) were ground using mortar and pestle under liquid nitrogen and pooled into a single sample per plant and stored at -80°C.

### RNA extraction and RNA-seq library preparation

Total RNA was extracted from each sample using the Spectrum Plant Total RNA kit (Sigma-Aldrich) following the manufacturer’s instructions and including DNAse treatment. Three samples per vineyard were generated by pooling 2 µg of total RNA from three plants from the same row in the vineyard for a total of 66 pools. Ribosome was depleted in 6 µg of RNA from each pool using the Dynabeads mRNA purification kit (Ambion, Invitrogen) following the manufacturer’s instructions. Ribosomal depleted RNA (25 ng per pool) was used as input for library preparation using the NEBNext Ultra RNA Library Prep Kit for Illumina (New England Biolabs Inc). Libraries were sequenced using Illumina NextSeq High Output 75 bp pair-end (Illumina Inc., San Diego, CA, United States) at the Australian Genome Research Facility (Adelaide, SA, Australia). Reads were trimmed and filtered using AdapterRemoval v2.2.1^39^ using default settings. Alignment of trimmed reads was performed in Hisat2 v2.1.0^40^ with default settings using the grapevine genome reference IGGP_12X, obtained from Ensembl Plants 45. BAM files from samples from the same vineyard were merged, sorted and indexed using SAMtools v1.8^41^. Mapped reads were counted to genomic features using featureCounts v1.5.2^42^, with the minimum mapping quality score for a read to be assigned to a feature was set to 10. The merged, sorted and indexed BAM files were then directly input into the R environment in order to identify the dicistronic tRNA-mRNA transcripts.

### *In silico* detection of dicistronic tRNA-mRNA transcripts

DiRT (**Di**cistronic **R**NA **T**ranscripts) is a custom pipeline implemented in the R environment and source codes are available at GitHub (https://github.com/CharlotteSai/DiRT). While the pipeline was developed for analysing grapevine RNA-seq data, it can be adapted for use in other species provided a genomic tRNA annotation is available. Firstly, protein coding gene (PCG) information and coordinates were downloaded from Ensembl Plants (release 45) (http://plants.ensembl.org/Vitis_vinifera/Info/Index) and the chromosomal coordinates of tRNA genes were extracted from the Genomic tRNA database using tRNAscan-SE based on predicted structure analysis (http://gtrnadb.ucsc.edu/GtRNAdb2/genomes/eukaryota/Vvini/). We used BEDTools version 2.25^43^ to determine the relative location of PCGs in relation to all tRNA genes (both upstream and downstream of the tRNA). Predicted tRNAs overlapping with PCGs were discarded for further analysis as there is no intergenic region between them and such reads could not be unambiguously assigned to either the tRNA or the PCG. tRNAs expressed (i.e. above 1 raw read) in leaf and berry samples were identified using GenomicRanges^44^. To infer putative co-transcription, first we filtered the RNA-seq data for genomic regions where both tRNA loci and closest neighbouring gene were transcribed (Raw read >= 1 for tRNAs and raw read >= 10 in PCGs), independently of their DNA-strand. A lower threshold of read coverage was applied for tRNAs due to underrepresentation in standard RNA-seq libraries^20^. Multi-mapped reads were used for mapping as there are over 600 predicted tRNA genes in *V. vinifera* often with multiple identifical isodecoder sequences which makes unique mapping of Illumina reads to individual transcribed tRNA loci challenging. In order to identify region-specific putative dicistronic transcripts, each Barossa sub-region was interrogated separately.

The selected candidate tRNA-mRNA transcripts were then scanned for dicistronic transcripts. We demanded that, first; the sequencing coverage of the intergenic region must be significantly higher than the intron closest to the intergenic region and the second intron closest to intergenic region. To achieve this, reads for each base of the intergenic region, the closest intron and the second closest intron were counted by the coverage method from the *GenomicRanges* package^44^ using merged BAM files for each region to obtain total coverage for each region. Then significant differences in average coverage between the intergenic region and the two closest introns were determined by a t-test including all the vineyards from the same sub-region as biological replicates (four vineyards for NG, CG, WR and EE, and three vineyards for SG and EV). If the protein coding gene contains just one intron, the t-test still is performed by comparing read coverage with the intergenic region. Of the 81 and 50 tRNA-mRNA pairs in leaf and berries respectively, 18 mRNAs contained one intron in both tissues and the remaining mRNAs contained two or more introns. The complete set of p-values were adjusted using the Benjamini-Hochberg false discovery rate (FDR)^45^ and intergenic regions with higher mean coverage than both introns, and an FDR-adjusted p-value < 0.05 were included for subsequent steps of the pipeline.

tRNA-mRNA transcripts passing the first condition were further filtered for those with uninterrupted sequencing coverage spanning the tRNA, the intergenic region and the mRNA by selecting candidates with at least one count for every base in the intergenic region. This condition was implemented to make sure that at least one entire molecule of the dicistronic transcript had been potentially produced.

Finally, dicistronic candidates with continuous coverage in the intergenic region were manually inspected using IGV^46^ for visual validation of continuous coverage. The candidates passing both of the t-tests and continuously coverage examination were deemed putative tRNA-mRNA dicistronic transcripts. Multidimensional scaling (MDS) were analysed in the R environment using the function *plotMDS* from the limma package^47^.

### RT-PCR confirmation

Complementary DNA was synthesized from the same total RNA used for the RNA-seq using SuperScript IV first strand synthesis system (Invitrogen, 18091050) following the manufacturer’s instructions. Complementary DNA was synthesised using gene specific reverse primers that aligned to the second exon of the candidate gene (for VIT_15s0046g02860_R, VIT_18s0001g09050_R), or the tRNA (tRNA^GlyCCC^_R) (+RT) and the reverse primer of the gene Elongation Factor 1-alpha (EF1a_R) as a positive control. Negative controls for the cDNA synthesis (-RT) in which reverse transcriptase enzyme was omitted were included for each of the dicistronic candidate. Resulting cDNA was diluted 1:10 and 2 µl was used for RT-PCR. The RT-PCR reaction was conducted using Kapa Taq PCR Kit (Kapa Biosystem, KK1020) following the manufacturer’s instructions. The amplification program used was 95°C for 3 min, 37 cycles at 95°C for 30 sec followed by 60°C for 30 sec and 72°C for 50 sec and finally 72°C per 2 min. For the candidate tRNA^ValCAC1.7^-VIT_15s0046g02860 we used primers tRNAVal^CAC^_F and Intergenic_tRNA^ValCAC^-VIT_15s0046g02860_R (376 bp). For the candidate tRNA^ProTGG2.9^-VIT_18s0001g09050 primers tRNA^ProTGG^_F and intergenic_tRNA^ProTGG^-VIT_18s0001g09050_R (172 bp) were used. For the candidate tRNA^GlyCCC^-VIT_19s0177g00220 we used primers VIT_19s0177g00220_F and intergenic_tRNAGlyCCC-VIT_19s0177g00220_R. Negative controls for the PCR reaction (-Ctr) contained all components for the reaction except the cDNA template. RT-PCR products were analysed by agarose gel electrophoresis and SYBR Safe DNA gel staining (ThermoFisher Scientific, S33102). RT-PCR products were purified using PCR Clean-up (Macherey-Nagel, 740609.250) following the manufacturer’s instructions. Sanger sequencing was performed at the Australian Genome Research Facility. Oligonucleotides used for RT-PCR are listed in Supplemental Table S6. Matching of the sequencing results for both putative dicistronic pairs and the expected sequence of each locus was confirmed using BLAST (blastn) with default settings^48^.

### Regional effect on the expression of dicistronic tRNA:mRNAs

To determine the effect of the region of origin on dicistronic tRNA:mRNAs, we independently compared the expression levels of all three components of the identified dicistronic transcripts (i.e. tRNAs, intergenic regions and PCGs). Similarly, the regional expression levels of all expressed tRNAs were compared. Briefly, mapped reads for each selected genomic feature obtained from featureCounts, were analysed in the R environment for plotting the gene expression through heatmaps. The expression of all tRNA, dicistronic tRNAs, intergenic region and the dicistronic genes (logCPM) were plotted using the pheatmap. Local Fisher LFDA was performed in the R environment using the package *lfda*^49^ to the expression (logCPM) values of all tRNA, dicistronic tRNAs, intergenic region and the dicistronic genes. In order to determine whether primary mRNA or the tRNA is driving the expression of candidate dicistronic tRNA-mRNA transcripts, we carried out Pearson correlation analyses between the expression of tRNA vs intergenic region and PCGs vs intergenic region for each dicistronic candidate identified. Pearson correlation analysis (p-value < 0.05) was performed using the R function *cor*.*test*(). Read counts of the intergenic regions were used as a proxy to define the expression of dicistronic transcripts. The rationale behind this lies in the assumption that reads mapping to the intergenic region can only be the result of the sequencing of a dicistronic RNA molecule, while reads mapping to tRNA genes and PCGs could result both from the expression of monocistronic and dicistronic transcripts (i.e. tRNA genes and PCGs pairs forming two independent RNA molecules or a single RNA molecules respectively) (Figure 6).

A bar plot was made to represent the distance (bp) between the tRNA and its proximal gene. Pairs of expressed tRNA-mRNA were split in two groups depending if they formed dicistronic or monocistronic transcripts. A non-overlapping sliding window approach (200bp) was used to count the number of pairs of genes of each type. Expression of the dicistronic genes and dicistronic tRNA was assessed by plotting their expression values against the distribution of the total gene expression for each tissue from the RNA-seq data. Gene annotation for dicistronic genes was obtained from the V1 annotation of the 12x grapevine reference genome (PN40024^18^), BLASTP search from NCBI (https://blast.ncbi.nlm.nih.gov/Blast.cgi), and from the Additional file 1 of Cramer et al.^50^. Protein information and gene ontology terms were obtained from UniProt (https://www.uniprot.org/uniprot). GO enrichment analysis was performed from Gene Ontology Consortium (http://geneontology.org/) with default settings.

### Motif analysis

Upstream and downstream sequence from the dicistronic tRNA was obtained from Genomic tRNA data and analysed in Weblogo^51^ for sequence analysis using default settings.

### Multi-species RNA-seq analysis

We collected short-read RNA sequencing datasets (Illumina paired and unpaired reads) from 30 independent studies (149 accessions) accessed through the National Centre for Biotechnology Information-Sequence Read Archive (NCBI-SRA) for nine land-plant species (i.e four vascular plants (*Arabidopsis thaliana, Vitis Vinifera, Oryza sativa* and *Brachypodium distachyon*), and three species with archaic vasculatures (*Azolla filiculoides, Salvinia cucullata* (ferns) and (*Selaginella moellendorffii* (lycophytes), and two non-vascular plants (*Physcomitrella patens* (mosses) and *Marchantia polymorpha* (liverworts)). See Supplemental Table S7 for library metadata and counts. Datasets were filtered based on the sample description provided to the SRA to include only whole organ or tissues samples that were likely to included vasculature. For species such as *Marchantia polymorpha* that do not contain true vasculature we included tissues that contain primitive vascular-like structures such as thallus, antheridiophore & archegoniophore. A minimum of three replicates per sample as denoted by the SRA metadata information was obtained. In order to reduce the experimental variables for the multi-species comparison, only wild-type plants and non-treated controls were included for downstream detection of tRNA-mRNA transcripts. The datasets were analysed using the DiRT pipeline as described previously.

### Expression analysis of orthologous genes

Identification of orthologous vascular genes in the lignin, cellulose and xylan pathways utilised a multi-step process. First, an OrthoFinder analysis was carried out as previously described^52^. The OrthoFinder algorithm^53^ generates orthogroups and subsequently infers gene trees for each orthogroup. To identify relevant orthogroups we then searched the orthofinder database for genes of interest (GOI) using gene codes from *Arabidopsis thaliana* (Supplemental Tables S8-10). Gene trees were reviewed by eye to identify orthologs of specific GOI. To ensure that orthologs were identified appropriately we then performed reciprocal BLAST searches for each candidate to confirm that orthologs were assigned appropriately. In some instances, it was challenging to unequivocally identify a single orthologous genes.

For the expression analysis of the identified orthologous genes, RNAseq datasets for the nine different plant species (Supplemental Table S11), were downloaded from NCBI’s Sequence Read Archive (SRA) database. Low quality reads were removed from the dataset using *trim_galore*.

Reads were then aligned to their respective species reference genome using HISAT2. After further processing of the aligned reads using the *samtools* package, read counts were calculated and normalized using TPMCalculator. The orthologous genes corresponding to the groups cellulose, xylan and lignin were then filtered from the dataset and their normalized expression was compared.

## Acknowledgments

This study was funded through a Pilot Program in Genomic Applications in Agriculture and Environment Sectors jointly supported by the University of Adelaide and the Australian Genome Research Facility Ltd. PJF was supported by Graduate Research Scholarships from Wine Australia (PH1503) and the University of Adelaide. NS was supported by a summer scholarship from the ARC Centre of Excellence in Plant Energy Biology (CE1400008). Dr Lopez is currently partially supported by the National Institute of Food and Agriculture, AFRI Competitive Grant Program Accession number 1018617 and National Institute of Food and Agriculture, United States Department of Agriculture, Hatch Program accession number 1020852. ERL wishes to acknowledge ongoing support from Prof Staffan Persson through ARC FT and DP funding (DP190101941; FT160100218). We thank the Barossa Grounds Project and the growers who allowed us to collect samples and supplied information about their vineyards. We thank Kendall Corbin for performing the DNA extraction of the leaf samples. We thank Cassandra Collins for the experimental design and collection of the plant material. We thank Roberta DeBei, Sandra Milena Mantilla, Annette James, and Valentin Olek who helped with the sample collection. We thank Stephen Tyerman for his contribution in the development of DiRT. We thank Timothy Cavagnaro and Andrew Metcalfe for their contribution in the experimental design. We thank Dr Uli Felzmann from Science IT, University of Melbourne, for assistance with high-performance computing infrastructure.

## Contributions

R. D. and C. M. R. L. conceived the project. P. T., J. B., M. G., C. M. R. L. and R. D. designed the research. P. J., L. A., A. S., B. C., F. Z. and R. D. performed the research. P. J., P. T. R. D. and C. M. R. L. wrote the paper. N. S. and S. P. developed the DiRT pipeline. All authors read and approved the final manuscript.

## Availability of data and materials

The full sequencing data are available on SRA database under the accession number PRJNA591273.

## References

1 Karginov TA, Pastor DPH, Semler BL, Gomez CM. Mammalian Polycistronic mRNAs and Disease. Trends Genet 2017; 33: 129–142.

2 Sugita M, Sugiura M. Regulation of gene expression in chloroplasts of higher plants. Plant Mol Biol 1996; 32: 315–326.

3 Barkan A. Expression of plastid genes: organelle-specific elaborations on a prokaryotic scaffold. Plant Physiol 2011; 155: 1520–1532.

4 Merchan F, Boualem A, Crespi M, Frugier F. Plant polycistronic precursors containing non-homologous microRNAs target transcripts encoding functionally related proteins. Genome Biol 2009; 10: R136.

5 Kruszka K, Barneche F, Guyot R, Ailhas J, Meneau I, Schiffer S et al. Plant dicistronic tRNA-snoRNA genes: a new mode of expression of the small nucleolar RNAs processed by RNase Z. EMBO J 2003; 22: 621–632.

6 Michaud M, Cognat V, Duchêne A-M, Maréchal-Drouard L. A global picture of tRNA genes in plant genomes. Plant J 2011; 66: 80–93.

7 Zhang W, Thieme CJ, Kollwig G, Apelt F, Yang L, Winter N et al. tRNA-related sequences trigger systemic mRNA transport in plants. Plant Cell 2016; 28: 1237–1249.

8 Banerjee R, Chen S, Dare K, Gilreath M, Praetorius-Ibba M, Raina M et al. tRNAs: cellular barcodes for amino acids. FEBS Lett 2010; 584: 387–395.

9 Dal Santo S, Tornielli GB, Zenoni S, Fasoli M, Farina L, Anesi A et al. The plasticity of the grapevine berry transcriptome. Genome Biol 2013; 14: r54.

10 Sun R, He F, Lan Y, Xing R, Liu R, Pan Q et al. Transcriptome comparison of Cabernet Sauvignon grape berries from two regions with distinct climate. J Plant Physiol 2015; 178: 43–54.

11 Sun X, Fan G, Su L, Wang W, Liang Z, Li S et al. Identification of cold-inducible microRNAs in grapevine. Front Plant Sci 2015; 6: 595.

12 Han J, Fang J, Wang C, Yin Y, Sun X, Leng X et al. Grapevine microRNAs responsive to exogenous gibberellin. BMC Genomics 2014; 15: 111.

13 Alabi OJ, Zheng Y, Jagadeeswaran G, Sunkar R, Naidu RA. High-throughput sequence analysis of small RNAs in grapevine (Vitis vinifera L.) affected by grapevine leafroll disease. Mol Plant Pathol 2012; 13: 1060–1076.

14 Wang C, Wang X, Kibet NK, Song C, Zhang C, Li X et al. Deep sequencing of grapevine flower and berry short RNA library for discovery of novel microRNAs and validation of precise sequences of grapevine microRNAs deposited in miRBase. Physiol Plant 2011; 143: 64–81.

15 Bester R, Burger JT, Maree HJ. Transcriptome analysis reveals differentially expressed small RNAs and genes associated with grapevine leafroll-associated virus 3 infections. Physiol Mol Plant Pathol 2017; 100: 220–236.

16 Yang Y, Mao L, Jittayasothorn Y, Kang Y, Jiao C, Fei Z et al. Messenger RNA exchange between scions and rootstocks in grafted grapevines. BMC Plant Biol 2015; 15: 251.

17 Robinson S, Sandercock N. An analysis of climate, soil and topographic information to aid the understanding of barossa terroir. SA: Primary Industries and Regions South Australia: Adelaide, Australia, 2014.

18 Jaillon O, Aury J-M, Noel B, Policriti A, Clepet C, Casagrande A et al. The grapevine genome sequence suggests ancestral hexaploidization in major angiosperm phyla. Nature 2007; 449: 463–467.

19 Chan PP, Lowe TM. GtRNAdb 2.0: an expanded database of transfer RNA genes identified in complete and draft genomes. Nucleic Acids Res 2016; 44: D184–9.

20 Burgess AL, David R, Searle IR. Conservation of tRNA and rRNA 5-methylcytosine in the kingdom Plantae. BMC Plant Biol 2015; 15: 199.

21 Yukawa Y, Sugita M, Choisne N, Small I, Sugiura M. The TATA motif, the CAA motif and the poly(T) transcription termination motif are all important for transcription re-initiation on plant tRNA genes. Plant J 2000; 22: 439–447.

22 Kersey PJ, Allen JE, Allot A, Barba M, Boddu S, Bolt BJ et al. Ensembl Genomes 2018: an integrated omics infrastructure for non-vertebrate species. Nucleic Acids Res 2018; 46: D802–D808.

23 Guan D, Yan B, Thieme C, Hua J, Zhu H, Boheler KR et al. PlaMoM: a comprehensive database compiles plant mobile macromolecules. Nucleic Acids Res 2017; 45: D1021–D1028.

24 Bowman JL, Kohchi T, Yamato KT, Jenkins J, Shu S, Ishizaki K et al. Insights into Land Plant Evolution Garnered from the Marchantia polymorpha Genome. Cell 2017; 171: 287–304.e15.

25 Ferrari C, Shivhare D, Hansen BO, Pasha A, Esteban E, Provart NJ et al. Expression Atlas of Selaginella moellendorffii Provides Insights into the Evolution of Vasculature, Secondary Metabolism, and Roots. Plant Cell 2020; 32: 853–870.

26 Thieme CJ, Rojas-Triana M, Stecyk E, Schudoma C, Zhang W, Yang L et al. Endogenous Arabidopsis messenger RNAs transported to distant tissues. Nat Plants 2015; 1: 15025.

27 Yang L, Perrera V, Saplaoura E, Apelt F, Bahin M, Kramdi A et al. m5C Methylation Guides Systemic Transport of Messenger RNA over Graft Junctions in Plants. Curr Biol 2019; 29: 2465–2476.e5.

28 Cui X, Liang Z, Shen L, Zhang Q, Bao S, Geng Y et al. 5-Methylcytosine RNA Methylation in Arabidopsis Thaliana. Mol Plant 2017; 10: 1387–1399.

29 David R, Burgess A, Parker B, Li J, Pulsford K, Sibbritt T et al. Transcriptome-Wide Mapping of RNA 5-Methylcytosine in Arabidopsis mRNAs and Noncoding RNAs. Plant Cell 2017; 29: 445–460.

30 Calderwood A, Kopriva S, Morris RJ. Transcript Abundance Explains mRNA Mobility Data in Arabidopsis thaliana. Plant Cell 2016; 28: 610–615.

31 Schramm L, Hernandez N. Recruitment of RNA polymerase III to its target promoters. Genes Dev 2002; 16: 2593–2620.

32 van Leeuwen C. Soils and Terroir Expression in Wines. In: Landa ER, Feller C (eds). Soil and Culture. Springer Netherlands: Dordrecht, 2009, pp 453–465.

33 Zsófi Z, Tóth E, Rusjan D, Bálo B. Terroir aspects of grape quality in a cool climate wine region: Relationship between water deficit, vegetative growth and berry sugar concentration. Sci Hortic (Amsterdam) 2011; 127: 494–499.

34 Gouthu S, O’Neil ST, Di Y, Ansarolia M, Megraw M, Deluc LG. A comparative study of ripening among berries of the grape cluster reveals an altered transcriptional programme and enhanced ripening rate in delayed berries. J Exp Bot 2014; 65: 5889–5902.

35 Rienth M, Torregrosa L, Luchaire N, Chatbanyong R, Lecourieux D, Kelly MT et al. Day and night heat stress trigger different transcriptomic responses in green and ripening grapevine (vitis vinifera) fruit. BMC Plant Biol 2014; 14: 108.

36 Ghaffari S, Reynard JS, Rienth M. Single berry reconstitution prior to RNA-sequencing reveals novel insights into transcriptomic remodeling by leafroll virus infections in grapevines. Sci Rep 2020; 10: 12905.

37 Xie H, Konate M, Sai N, Tesfamicael KG, Cavagnaro T, Gilliham M et al. Global DNA methylation patterns can play a role in defining terroir in grapevine (Vitis vinifera cv. Shiraz). Front Plant Sci 2017; 8: 1860.

38 Williams LE, Araujo FJ. Correlations among Predawn Leaf, Midday Leaf, and Midday Stem Water Potential and their Correlations with other Measures of Soil and Plant Water Status in Vitis vinifera. J Amer Soc Hort Sci 2002; 127: 448–454.

39 Schubert M, Lindgreen S, Orlando L. AdapterRemoval v2: rapid adapter trimming, identification, and read merging. BMC Res Notes 2016; 9: 88.

40 Kim D, Langmead B, Salzberg SL. HISAT: a fast spliced aligner with low memory requirements. Nat Methods 2015; 12: 357–360.

41 Li H, Handsaker B, Wysoker A, Fennell T, Ruan J, Homer N et al. The Sequence Alignment/Map format and SAMtools. Bioinformatics 2009; 25: 2078–2079.

42 Liao Y, Smyth GK, Shi W. featureCounts: an efficient general purpose program for assigning sequence reads to genomic features. Bioinformatics 2014; 30: 923–930.

43 Quinlan AR, Hall IM. BEDTools: a flexible suite of utilities for comparing genomic features. Bioinformatics 2010; 26: 841–842.

44 Lawrence M, Huber W, Pagès H, Aboyoun P, Carlson M, Gentleman R et al. Software for computing and annotating genomic ranges. PLoS Comput Biol 2013; 9: e1003118.

45 Benjamini Y, Hochberg Y. Controlling the False Discovery Rate: A Practical and Powerful Approach to Multiple Testing. J R Statist Soc 1995; 57: 239–300.

46 Robinson JT, Thorvaldsdóttir H, Winckler W, Guttman M, Lander ES, Getz G et al. Integrative genomics viewer. Nat Biotechnol 2011; 29: 24–26.

47 Ritchie MD, Holzinger ER, Li R, Pendergrass SA, Kim D. Methods of integrating data to uncover genotype-phenotype interactions. Nat Rev Genet 2015; 16: 85–97.

48 Zhang Z, Schwartz S, Wagner L, Miller W. A greedy algorithm for aligning DNA sequences. J Comput Biol 2000; 7: 203–214.

49 Sugiyama M. Local Fisher discriminant analysis for supervised dimensionality reduction. In: Proceedings of the 23rd international conference on Machine learning - ICML ’06. ACM Press: New York, New York, USA, 2006, pp 905–912.

50 Cramer GR, Cochetel N, Ghan R, Destrac-Irvine A, Delrot S. A sense of place: transcriptomics identifies environmental signatures in Cabernet Sauvignon berry skins in the late stages of ripening. BMC Plant Biol 2020; 20: 41.

51 Crooks GE, Hon G, Chandonia JM, Brenner SE. WebLogo: a sequence logo generator. Genome Res 2004; 14: 1188–1190.

52 Lampugnani ER, Flores-Sandoval E, Tan QW, Mutwil M, Bowman JL, Persson S. Cellulose Synthesis - Central Components and Their Evolutionary Relationships. Trends Plant Sci 2019; 24: 402–412.

53 Emms DM, Kelly S. OrthoFinder: solving fundamental biases in whole genome comparisons dramatically improves orthogroup inference accuracy. Genome Biol 2015; 16: 157.

