## Supplemental Fig1 for "Tissue and regional expression patterns of dicistronic tRNA-mRNA transcripts in grapevine (*Vitis vinifera*) and their evolutionary co-appearance with vasculature in land plants"

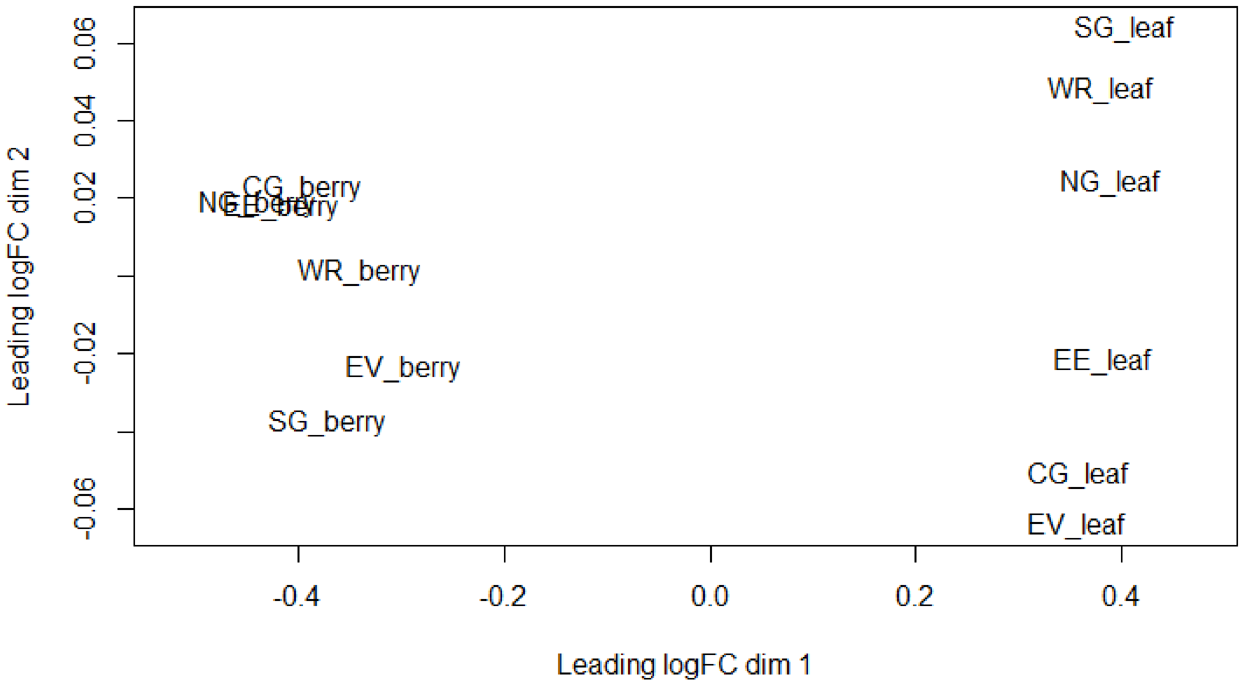


Supplemental_Fig_S1.pdf: MDS plots of total raw counts of all tRNAs expressed in leaf and berry samples for each sub-region.
