## Supplemental Fig2 for "Tissue and regional expression patterns of dicistronic tRNA-mRNA transcripts in grapevine (*Vitis vinifera*) and their evolutionary co-appearance with vasculature in land plants"

a)

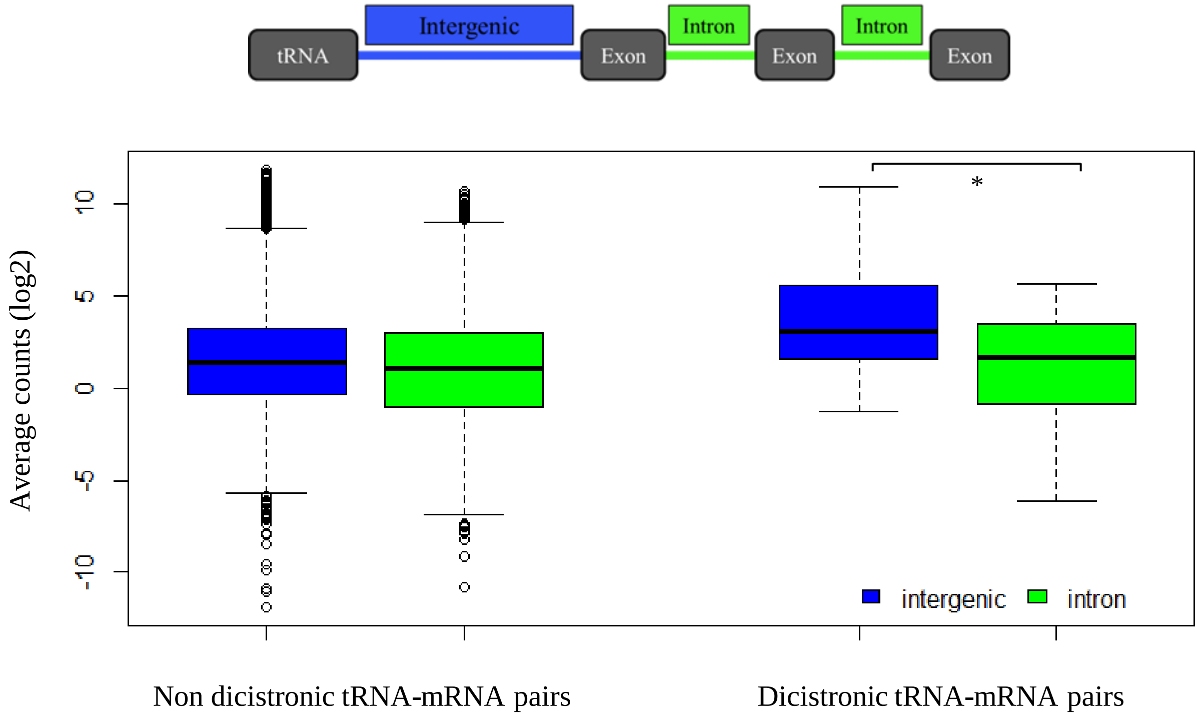


b)

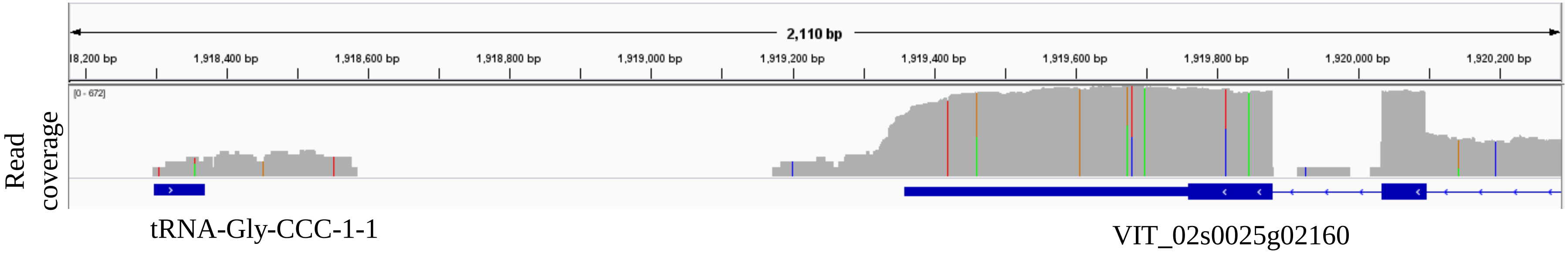


Supplemental_Fig_S2.pdf: Comparative analysis of expression levels of tRNA-mRNA intergenic regions (blue) and closest two introns (green) in non-dicistronic (left) and putative dicistronic tRNA-mRNA pairs (right) in leaf tissue. a) Average expression of intergenic region was significantly higher than the two closest introns in dicistronic tRNA-mRNA pairs (t-test, p-value $< 0.05$ ). b) Genome browser view of a non-dicistronic tRNA-mRNA pair (tRNA-Gly-CCC-1-1_VIT_02s0025g02160) identified by the DiRT pipeline. In this example, the tRNA and PCG are both expressed but the intergenic region does not present continuous coverage.
