## Supplemental Fig3 for "Tissue and regional expression patterns of dicistronic tRNA-mRNA transcripts in grapevine (*Vitis vinifera*) and their evolutionary co-appearance with vasculature in land plants"

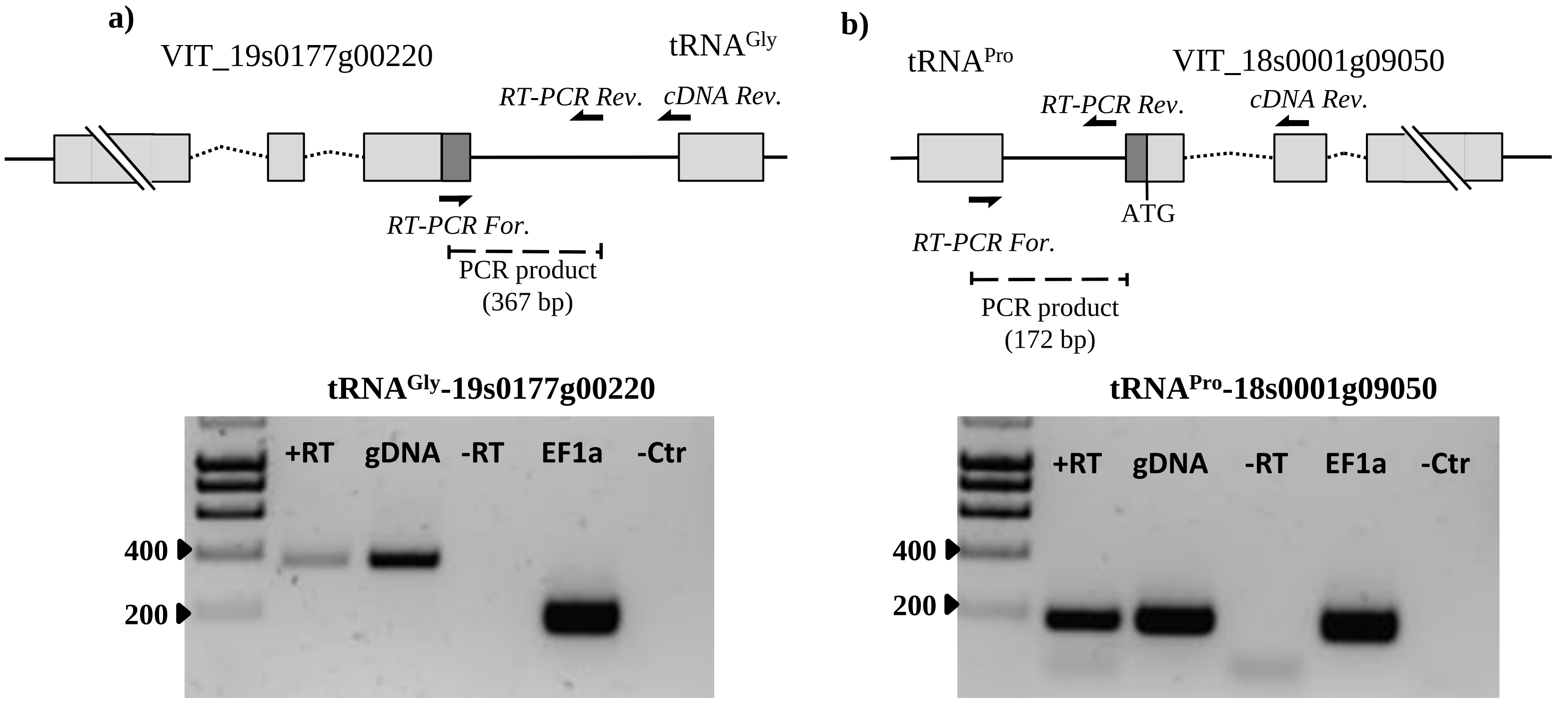


Supplemental_Fig_S3.pdf: RT-PCR confirmation of identified dicistronic transcripts in berry samples. Model of putative dicistronic tRNA-mRNA transcript showing primers used for cDNA synthesis (cDNA rev.) and for the PCR reaction (RT-PCR For and RT-PCR Rev). Confirmation of actively transcribed intergenic region through RT-PCR for candidates a) $tRNA^{GlyCCC}$_VIT_19s0177g00220 (367 bp) and b) $tRNA^{ProTGG}$-VIT_18s0001g09050 (172 bp). +RT: cDNA as template, gDNA: genomic DNA was used as a control, -RT: RT-PCR negative control, EF1a: Elongation Factor 1-alpha was used as a positive control (150 bp), -Ctr: PCR negative control.
