## Supplemental Fig4 for "Tissue and regional expression patterns of dicistronic tRNA-mRNA transcripts in grapevine (*Vitis vinifera*) and their evolutionary co-appearance with vasculature in land plants"

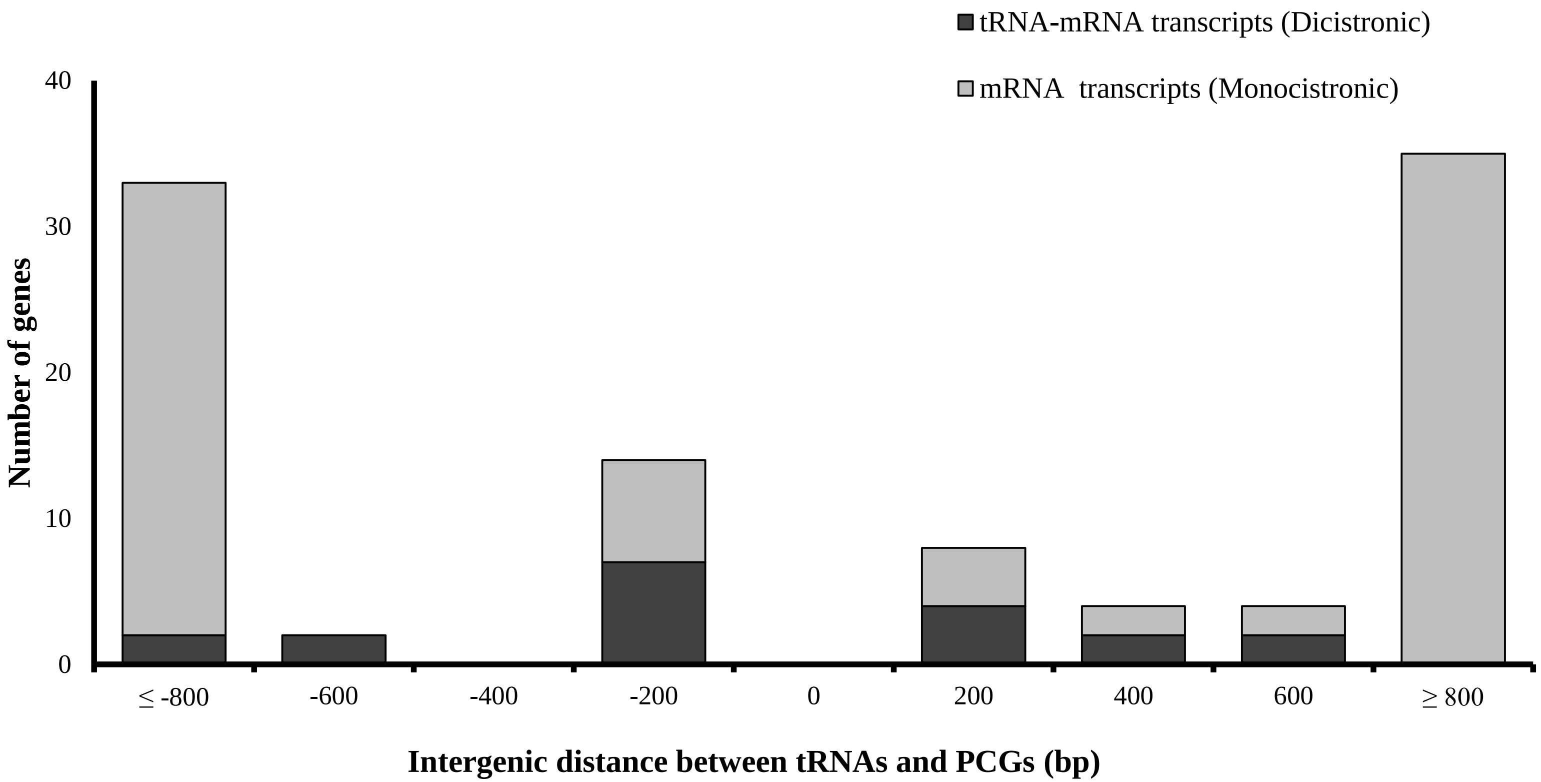


Supplemental_Fig_S4.pdf: Frequency of the length of the intergenic distance between tRNA (centred at zero) and protein coding genes (PGCs) for dicistronic and monocistronic transcripts. Vertical bars show the number of neighbouring tRNAs and PCGs pairs forming putative dicistronic tRNA-mRNA transcripts (black), or monocistronic transcripts (grey).
