## Supplemental Fig5 for "Tissue and regional expression patterns of dicistronic tRNA-mRNA transcripts in grapevine (*Vitis vinifera*) and their evolutionary co-appearance with vasculature in land plants"

**a)**

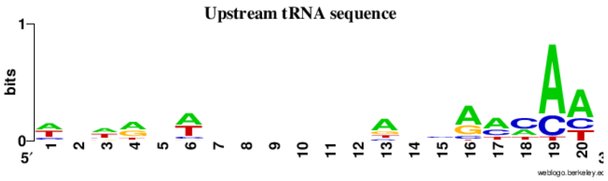


**b)**

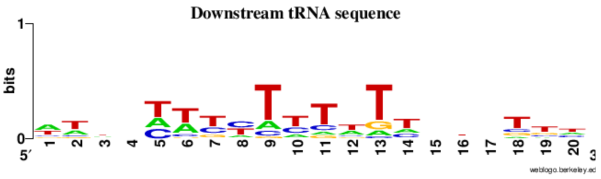


Supplemental_Fig_S5.pdf: Weblogo sequence analysis of the first 20 bp a) upstream; b) downstream of the 19 candidates dicistronic tRNAs.
