## Supplemental Fig6 for "Tissue and regional expression patterns of dicistronic tRNA-mRNA transcripts in grapevine (*Vitis vinifera*) and their evolutionary co-appearance with vasculature in land plants"

Leaf

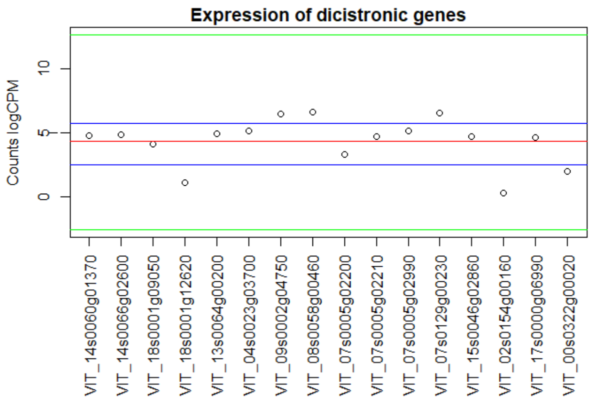


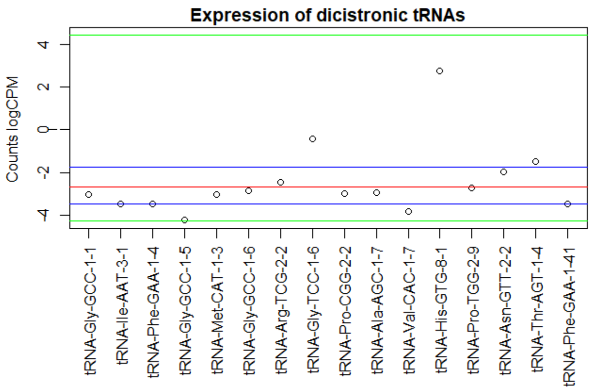


Berry

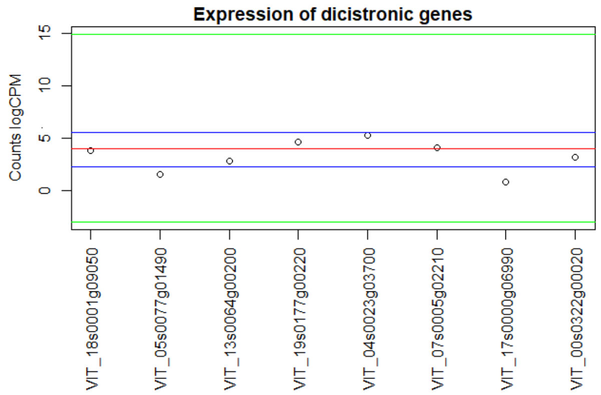


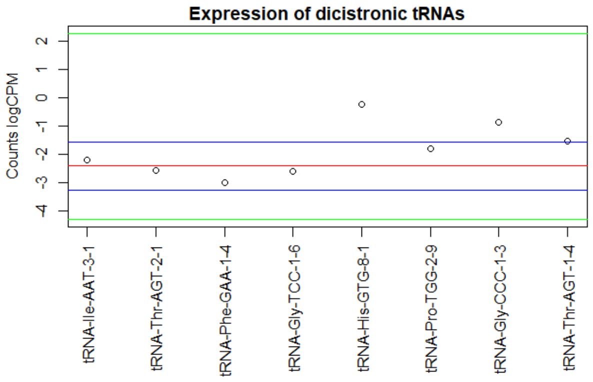


Supplemental_Fig_S6.pdf: Expression of the candidate dicistronic genes and tRNA with respect of the total gene and tRNAs expressed from the RNA-seq data for both tissues. Expression of candidates dicistronic genes compared with the expression of 18,698 genes in leaf and 17,160 genes in berry samples. Expression of candidates dicistronic tRNAs were compared against the expression of total tRNA expressed in leaf (124) and berry (90) samples. Red line represent the median of the total expression. Blue lines correspond to the 25% and 75% quartile of the distribution of the total gene expression. Green lines are the minimum and maximum of gene expression.
