## Supplemental Fig7 for "Tissue and regional expression patterns of dicistronic tRNA-mRNA transcripts in grapevine (*Vitis vinifera*) and their evolutionary co-appearance with vasculature in land plants"

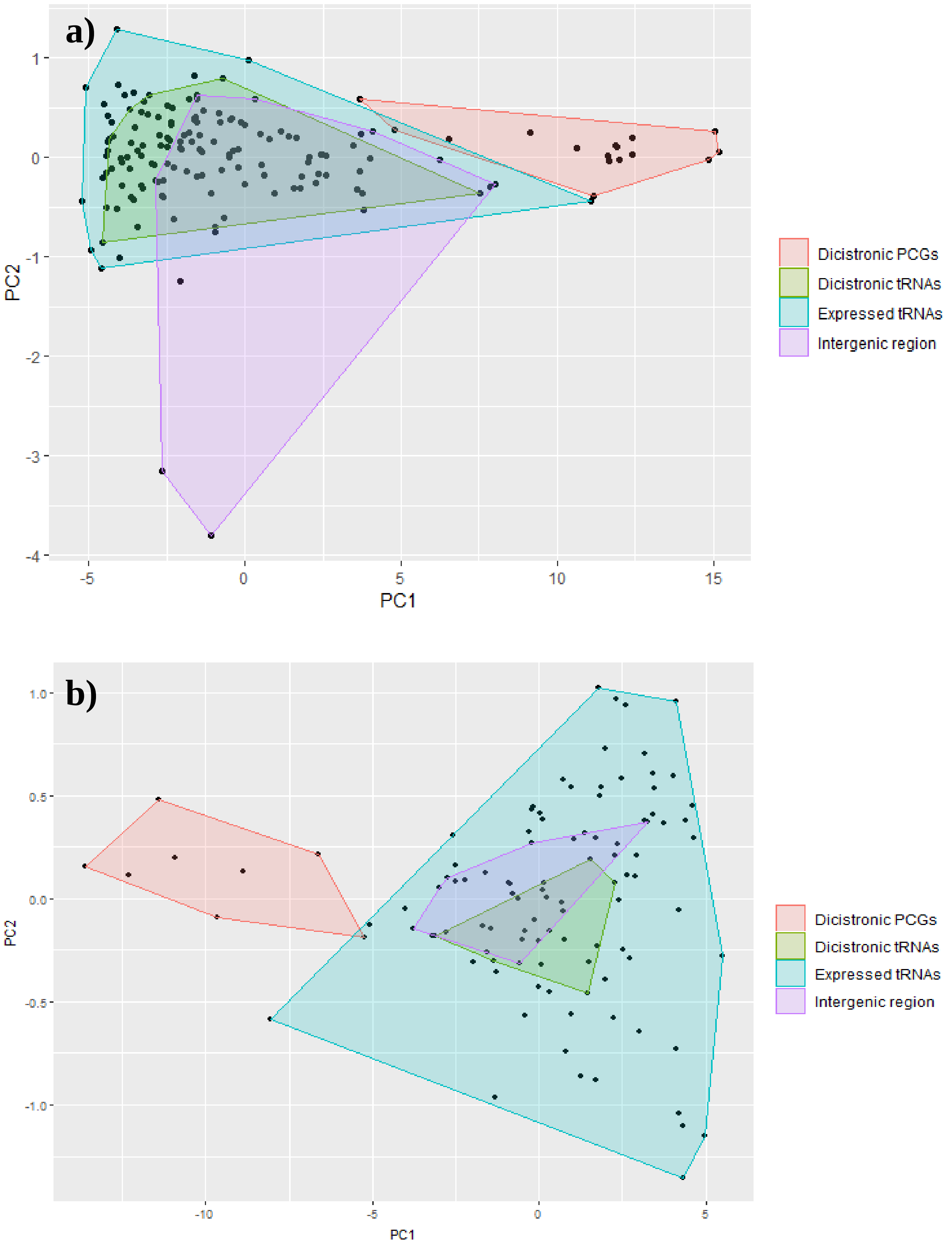


Supplemental_Fig_S7.pdf: Local Fisher Discriminant Analysis of the expression (logCPM) of all expressed tRNAS, dicistronic tRNAs, intergenic region and dicistronic genes for each sub-region from the Barossa Wine growing region for a) leaf and b) berry samples. Clustering of the samples allows to clearly differentiate the expression of the PCGs from the tRNAs and intergenic regions.
