## Supplemental Fig8 for "Tissue and regional expression patterns of dicistronic tRNA-mRNA transcripts in grapevine (*Vitis vinifera*) and their evolutionary co-appearance with vasculature in land plants"

a)

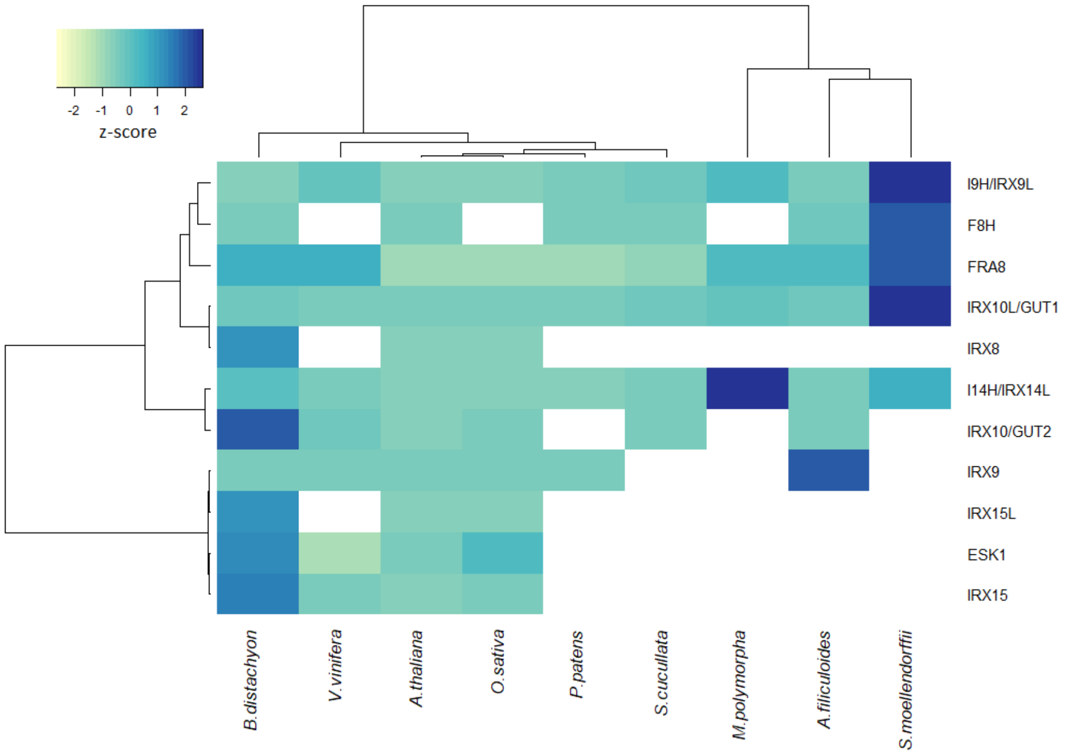


b)

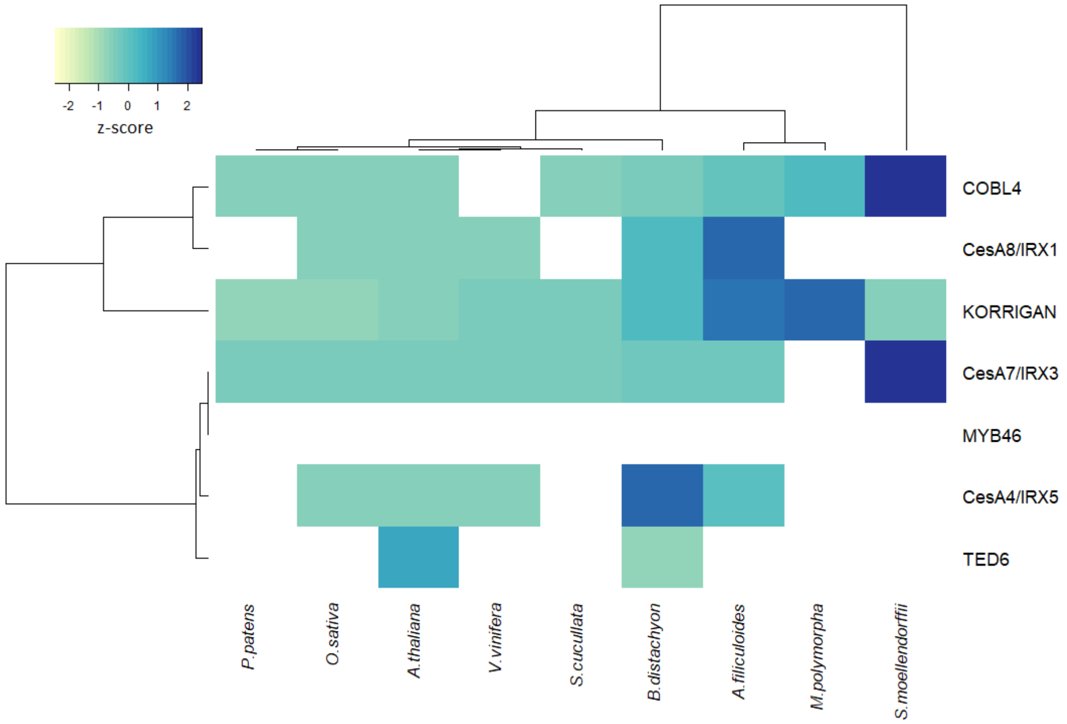


Supplemental Figure S: RNA-seq expression heat map (TPM) of orthologous xylan and cellulose biosynthesis genes involved in vasculature development. Average TPM expression values were derived from the same RNA-seq datasets used for analysing dicistronic tRNA-mRNA transcription. Colour represents the expression Z-score (TPM minus mean over s.d) of each gene (row) in a given species (column). A white panel is used to indicate cases where unequivocal ortholog assignment was not possible or wherean orthologous gene member could not be identified.
