## Supplemental Table1 for "Tissue and regional expression patterns of dicistronic tRNA-mRNA transcripts in grapevine (*Vitis vinifera*) and their evolutionary co-appearance with vasculature in land plants"

**Supplementary Table S1**: Mapped RNA-seq data for leaf and berry samples to the *V. vinifera* to the reference genome PN40024.

| Samples | Reads leaf samples | Reads berry samples | Merging, sorting and indexing of samples | Vineyard | Reads per vineyard leaf samples | Reads per vineyard berry samples |
|---|---|---|---|---|---|---|
| CG1.1 | 25,743,097 | 25,531,723 | | | | |
| CG1.2 | 27,225,555 | 24,411,207 | | | | |
| CG1.3 | 26,383,747 | 23,687,732 | | CG1 | 79,352,399 | 73,630,662 |
| CG2.1 | 20,726,878 | 23,885,740 | | | | |
| CG2.2 | 37,326,602 | 26,059,204 | | | | |
| CG2.3 | 20,330,612 | 25,424,851 | | CG2 | 78,384,092 | 75,369,795 |
| CG3.1 | 25,863,809 | 24,002,370 | | | | |
| CG3.2 | 20,856,760 | 20,203,746 | | | | |
| CG3.3 | 19,566,185 | 18,694,547 | | CG3 | 66,286,754 | 62,900,663 |
| CG4.1 | 25,342,050 | 20,911,435 | | | | |
| CG4.2 | 22,221,312 | 22,605,405 | | | | |
| CG4.3 | 22,097,859 | 26,006,395 | | CG4 | 69,661,221 | 69,523,235 |
| EV1.1 | 23,056,320 | 20,946,939 | | | | |
| EV1.2 | 25,524,736 | 23,149,751 | | | | |
| EV1.3 | 17,605,976 | 23,905,651 | | EV1 | 66,187,032 | 68,002,341 |
| EV2.1 | 25,353,854 | 16,541,723 | | | | |
| EV2.2 | 13,559,924 | 16,291,835 | | | | |
| EV2.3 | 21,162,499 | 21,175,449 | | EV2 | 60,076,277 | 54,009,007 |
| EV3.1 | 10,817,673 | 20,664,463 | | | | |
| EV3.2 | 21,138,475 | 22,523,299 | | | | |
| EV3.3 | 25,676,349 | 22,571,290 | | EV3 | 57,632,497 | 65,759,052 |
| NG1.1 | 29,778,866 | 20,227,972 | | | | |
| NG1.2 | 24,798,843 | 18,273,130 | | | | |
| NG1.3 | 54,236,656 | 22,558,469 | | NG1 | 108,814,365 | 61,059,571 |
| NG2.1 | 24,638,270 | 19,482,965 | | | | |
| NG2.2 | 29,024,880 | 16,554,581 | | | | |
| NG2.3 | 18,360,950 | 21,490,872 | | NG2 | 72,024,100 | 57,528,418 |
| NG3.1 | 32,656,908 | 18,957,727 | | | | |
| NG3.2 | 15,254,356 | 18,703,259 | | | | |
| NG3.3 | 21,842,503 | 22,475,849 | | NG3 | 69,753,767 | 60,136,835 |
| NG4.1 | 18,310,992 | 23,146,019 | | | | |
| NG4.2 | 27,310,080 | 29,280,200 | | | | |
| NG4.3 | 19,468,484 | 19,077,555 | | NG4 | 65,089,556 | 71,503,774 |
| EE1.1 | 31,012,663 | 18,810,900 | | | | |
| EE1.2 | 30,189,945 | 16,909,798 | | | | |
| EE1.3 | 26,985,753 | 20,316,791 | | EE1 | 88,188,361 | 56,037,489 |
| EE2.1 | 42,939,482 | 19,603,339 | | | | |
| EE2.2 | 29,521,962 | 20,177,138 | | | | |

| | | | | | |
|---|---|---|---|---|---|
| EE2.3 | 25,145,787 | 21,774,682 | EE2 | 97,607,231 | 61,555,159 |
| EE3.1 | 29,216,096 | 22,752,201 | | | |
| EE3.2 | 18,263,688 | 21,445,703 | | | |
| EE3.3 | 21,728,373 | 20,282,356 | EE3 | 69,208,157 | 64,480,260 |
| EE4.1 | 11,557,040 | 19,026,973 | | | |
| EE4.2 | 13,484,751 | 16,911,470 | | | |
| EE4.3 | 8,873,232 | 18,423,419 | EE4 | 33,915,023 | 54,361,862 |
| SG1.1 | 15,484,201 | 21,718,901 | | | |
| SG1.2 | 17,223,812 | 19,702,658 | | | |
| SG1.3 | 17,162,388 | 17,652,784 | SG1 | 49,870,401 | 59,074,343 |
| SG2.1 | 16,276,961 | 18,821,186 | | | |
| SG2.2 | 18,392,528 | 22,273,698 | | | |
| SG2.3 | 16,084,122 | 21,023,802 | SG2 | 50,753,611 | 62,118,686 |
| SG3.1 | 14,614,325 | 16,408,686 | | | |
| SG3.2 | 17,965,039 | 20,178,538 | | | |
| SG3.3 | 23,806,350 | 20,694,700 | SG3 | 56,385,714 | 57,281,924 |
| WR1.1 | 16,338,124 | 21,886,857 | | | |
| WR1.2 | 20,531,631 | 21,246,529 | | | |
| WR1.3 | 42,524,626 | 24,688,790 | WR1 | 79,394,381 | 67,822,176 |
| WR2.1 | 21,867,390 | 21,933,643 | | | |
| WR2.2 | 21,709,202 | 27,810,725 | | | |
| WR2.3 | 17,494,025 | 22,295,422 | WR2 | 61,070,617 | 72,039,790 |
| WR3.1 | 21,713,180 | 23,717,426 | | | |
| WR3.2 | 18,808,425 | 19,326,702 | | | |
| WR3.3 | 23,660,660 | 24,110,068 | WR3 | 64,182,265 | 67,154,196 |
| WR4.1 | 20,540,628 | 21,293,512 | | | |
| WR4.2 | 44,712,982 | 23,626,566 | | | |
| WR4.3 | 17,825,928 | 24,641,608 | WR4 | 83,079,538 | 69,561,686 |
