## Supplemental Table2 for "Tissue and regional expression patterns of dicistronic tRNA-mRNA transcripts in grapevine (*Vitis vinifera*) and their evolutionary co-appearance with vasculature in land plants"

Supplementary table S2: Percentage of each tRNA isotype expressed in leaf and berry samples.

| Isotype | Leaf % | Berry % |
|---|---:|---:|
| Pro | 16% | 18% |
| Ala | 9% | 9% |
| Arg | 8% | 8% |
| Gly | 8% | 6% |
| Gln | 6% | 4% |
| Asn | 5% | 4% |
| Thr | 5% | 7% |
| Met | 5% | 4% |
| Val | 5% | 6% |
| Leu | 5% | 2% |
| His | 4% | 3% |
| Ile | 4% | 4% |
| Cys | 3% | 4% |
| Phe | 3% | 3% |
| Tyr | 3% | 3% |
| Ser | 3% | 4% |
| Glu | 3% | 4% |
| Asp | 2% | 1% |
| Lys | 1% | 1% |
| Trp | 0% | 1% |
