## Supplemental Table3 for "Tissue and regional expression patterns of dicistronic tRNA-mRNA transcripts in grapevine (*Vitis vinifera*) and their evolutionary co-appearance with vasculature in land plants"

| **Supplementary table S3**: Sanger sequencing result of the RT-PCR product from the intrergenic region of two dicistronic candidates (cDNA) | | | PCR product were align to the query sequence in Blast2sequence with defult settings | | | | | | | | |
|---|---|---|---|---|---|---|---|---|---|---|---|
| Name | primer | Sequencing results | Query sequence: tRNA + intergenic region | Target ID | seq primer | Sequence length | Max Score | Total Score | Query Cover | E value | Per. Ident |
| $tRNA^{ProTGG}$ | Forward | >1_ProC_F_B09<br>TTTCTTTGGCTGGGTTTTGGTTTTACTTCACC<br>ATAAACCTCAAAAAAGCCCTCTTATGCT<br>CTTCTGCAAATTTCATTTGTGTTATTGGTACT<br>GAAACTCCGAGGCGGTGGCAGGCAAGGA<br>AACAG | ">tRNA-Pro-TGG-2-9_Intergenic_region<br>TGCGAGAGGTCCCGAGTTCGATT<br>CTCGGAATGCCCCAATCTTTTTATTTTCTTTCGCTGGGTTT<br>TGGTTTTACTTCACCATAA<br>ACCTCAAAAAAGCCCTCTTATGCTCTTCTGCAAATTTCAT<br>TTGTGTTATTGGTACTGAAA<br>CTCCGAGGCGGTGGCAGGCAAGGAAACA" | tRNA-Pro-TGG-2-9_Intergenic_region | Forward (F) | 172 | 224 | 224 | 0.72 | 9E-64 | 99.2% |
| Intergenic_$tRNA^{ProTGG}$-Vv18s0001g09 | Reverse | >2_ProC_R_B10<br>TGAATTTGCAGAGAGCATAAGAGGGCTTTTT<br>TGAGGTTTATGGTGAAGTAAAACCAAAAC<br>CCAGCGAAAGAAAATAAAAAGATTGGGGCA<br>TTCCGAGAATCGAACTCGGGACCTCTCGCA | CGGGCATTTGGTCTAGTGGTATGATTCTCGCTTTGGGTGC<br>GAGAGGTCCCGAGTTCGATT<br>CTCGGAATGCCCCAATCTTTTTATTTTCTTTCGCTGGGTTT<br>TGGTTTTACTTCACCATAAACCTCAAAAAAGC<br>CCTCTTATGCTCTTCTGCAAATTTCATTTGTGTTATTGGTA<br>CTGAAACTCCGAGGCGGTG<br>GCAGGCAAGGAAA | | Reverse (R ) | | 213 | 213 | 0.69 | 6E-60 | 99.2% |
| $tRNA^{ValCAC}$ | Forward | >5_ValC_F_C01<br>ATTGCCAC-<br>AGTCTTCCATTTCTGTTGGGAGTCTCCCAGGG<br>TCAGAGTATCAACGACACT<br>CAGTGCCACAATCTCATTTCCATTTCTGCTAG<br>GAGTCTCCCAGATTCTCAATATCAACAA<br>CACTCACCCAGATTTTTAACATTTTCTCATCT<br>GGATGTTCATCAATTAGTCAAACAATGC<br>AGATTCAGCCACACCCACTTCCAAACTATAGT<br>CTTAGGTCACCAATTTTCTCACCTTCGA<br>CCCATTTTCTCATACCTTTCTCATCATCATTCC<br>AACCTACTAAACTTGTCTCTATTCCAA<br>ATTTCCCATCTGGGTCTTGCG | ">tRNA-Val-CAC-1-7_Intergenic_region<br>CACTAGAGGTCCCCGGTTCGAACCCGGGCTCAGACA<br>TTTGCATTTTTATTTTATTATTGCCAGAGTCTTCCATTTCT<br>GTTG<br>GGAGTCTCCCAGGGTCAGAGTATCAACGACACTCAGTGC<br>CACAATCTCATTTCCATTTCT<br>GCTAGGAGTCTCCCAGATTCTCAATATCAACAACACTCAC<br>CCAGATTTTTAACATTTTCT<br>CATCTGGATGTTCATCAATTAGTCAAACAATGCAGATTCA<br>GCCACACCCACTTCCAAACT<br>ATAGTCTTAGGTCACCAATTTTCTCACCTTCGACCCATTTT<br>CTCATACCTTTCTCATCAT<br>CATTCCAACCTACTAAACTTGTCTCTATTCCAAATTTCCCA<br>TCTGGGTCTTGCGG" | | Forward (F) | 376 | 586 | 586 | 0.85 | 6E-172 | 99.7% |
| Intergenic_$tRNA^{ValCAC}$-Vv15s0046g02860 | Reverse | >6_ValC_R_C02<br>ATGAGAAAATGGGTCGAAGGTGAGAAAATT<br>GGTGACCTAAGACTATAGTTTGGAAGTGGG<br>TGTGGCTGAATCTGCATTGTTTGACTAATTGA<br>TGAACATCCAGATGAGAAAATGTTAAAA<br>ATCTGGGTGAGTGTTGTTGATATTGAGAATC<br>TGGGAGACTCCTAGCAGAAATGGAAATGA<br>GATTGTGGCACTGAGTGTCGTTGATACTCTG<br>ACCCTGGGAGACTCCCAACAGAAATGGAA<br>GACTGTGGCAATAATAAAATAAAAATGCAAA<br>TGTCTGAGCCCGGGTTCGAACCGGGGACC<br>TCTAGTG | | tRNA-Val-CAC-1-7_Intergenic_region | Reverse (R ) | | 562 | 562 | 0.81 | 1E-164 | 99.7% |
