## Supplemental Table4 for "Tissue and regional expression patterns of dicistronic tRNA-mRNA transcripts in grapevine (*Vitis vinifera*) and their evolutionary co-appearance with vasculature in land plants"

**Supplementary Table S4**: Gene ontology terms and annotated function of dicistronic genes identified in grapevine leaf, berry and common between both tissues. Function of the genes was determined by gene ontology motif that the genes presented (BLASTP) and from Cramer et al. (2020). Closest Arabidopsis match to the grapevine PCG candidates (based on protein sequence BLAST search) and evidence of dicistronic transcription and mobility.

| **Gene ID** | **RefSeq** | **Gene function** | **GO terms** | **Description of the GO term** | **BLAST top *Arabidopsis* match (Ensembl BLAST results)** | **Description *A. thaliana* (source Araport11)** | **Dicistronic in *Arabidopsis* (Zhang et al. 2016)** | **Common dicistronic tRNA in Arabidopsis and grapevine** | **mRNA mobile in Arabidopsis (PlaMoM database)** | **Experimental evidence for mobility (PubMed ID)** |
|---|---|---|---|---|---|---|---|---|---|---|
| VIT_09s0002g04750 | XP_010654677.1 | MAR-binding filament-like protein 1-1 isoform | GO:0006259<br>GO:0005622<br>GO:0006950 | DNA metabolic process<br>Intracellular<br>Response to stress | AT1G05320 (19.4% AA ID, 0.0011 E-val) | PICC-LIKE, PICL myosin heavy chain, embryonic smooth protein | No | - | No | - |
| VIT_08s0058g00460 | XP_002263502.1 | Uncharacterized protein | No GO terms | No GO terms | AT1G02816 (27.8% AA ID, 2.7E-8 E-val) | pectinesterase | No | - | No | - |
| VIT_07s0129g00230 | XP_002279597.1 | Glucan endo-1,3-beta-glucosidase 1 | GO:0004553<br>GO:0005975 | hydrolase activity, hydrolyzing O-glycosyl compounds<br>carbohydrate metabolic process | AT5G67460 (76.9% AA ID, 1.1E-48 E-val) | O-Glycosyl hydrolases family 17 protein | Yes | tRNA-Met(CAT) | No | - |
| VIT_14s0060g01370 | XP_019080365.1 | Zinc finger matrin-type protein 2 isoform | GO:0003676<br>GO:0008270 | nucleic acid binding<br>zinc ion binding | AT3G05760 (87.9% AA ID, 5.0E-62 | C2H2 and C2HC zinc fingers superfamily protein | No | - | Yes | EDTA-exudation (1848506 10 |

| | | | | | | | | | | |
|---|---|---|---|---|---|---|---|---|---|---|
| | | | GO:0000398 | mRNA splicing, via spliceosome | E-val) | | | | | |
| VIT_18s0001g12620 | XP_010664706.1 | RING-type E3 ubiquitin transferase | GO:0004842 | ubiquitin-protein transferase activity | AT1G29340 (45.9% AA ID, 3.5E-21 E-val) | ATPUB17 Encodes a protein containing a UND, a U-box, and an ARM domain | No | - | Yes | grafting (27247031) |
| VIT_02s0154g00160 | XP_010662302.1 | Flavin-containing monooxygenase FMO GS-OX-like 3 | GO:0050660 | flavin adenine dinucleotide binding | AT1G62540 (60.6% AA ID, 1.7E-39 E-val) | belongs to the flavin-monooxygenase (FMO) family, encodes a glucosinolate S-oxygenase that catalyzes the conversion of methylthioalkyl glucosinolates to methylsulfinylalkyl glucosinolates | No | - | Yes | grafting (27247031) |
| | | | GO:0004497 | monooxygenase activity | | | | | | |
| | | | GO:0004499 | N,N-dimethylaniline monooxygenase activity | | | | | | |
| | | | GO:0050661 | NADP binding | | | | | | |
| VIT_07s0005g02200 | XP_002280035.1 | Autophagy-related 8i-like | GO:0006914 | autophagy | AT3G15580 (78.3% AA ID, 3.8E-59 E-val) | APG8H, Encodes APG8, a component of autophagy conjugation pathway. Delivered to the lumens of vacuole under nitrogen-starvation condition. | Yes | tRNA-Phe(GAA) | Yes | LMPC-exudation, EDTA-exudation (18485061) |
| | | | GO:0006995 | cellular response to nitrogen starvation | | | | | | |
| VIT_15s0046g02860 | XP_002273879.2 | Riboflavin biosynthesis protein PYRD, chloroplastic | GO:0008835 | diaminohydroxyphosphoribosylaminopyrimidine deaminase activity | AT3G47390 (36% AA ID, 2.3E-4 E-val) | PHOTOSENSITIVE 1 Encodes a protein that is believed to function as a pyrimidine reductase involved | No | - | No | - |
| | | | GO:0008270 | zinc ion binding | | | | | | |

| | | | | | | | | | | |
|---|---|---|---|---|---|---|---|---|---|---|
| | | | GO:0055114 | oxidation-reduction process | | in riboflavin and FAD biosynthesis. phs1 was identified as a photosensitive mutant that shows reduced growth, chloroplast developmental abnormalities, reduced chlorophyll levels, increased oxidative stress, reduced NADPH/NADP+ ratios, reduced photosystem I electron transport, reduced chlorophyll levels, increased oxidative stress, reduced NADPH/NADP+ ratios, reduced photosystem I electron transport, and reduced photosynthetic protein levels under high light conditions. Many of these abnormal phenotypes likely arise from the reduction in the levels of FAD in the phs1 mutant. | | | | |
| | | | GO:0009231 | riboflavin biosynthetic process | | | | | | |
| VIT_07s0005g02990 | XP_002262985.1 | Bet1-like SNARE 1-1 | GO:0005484 | SNAP receptor activity | AT3G58170 (75% AA ID, 4.7E-48 E-val) | ARABIDOPSIS THALIANA BET1P/SFT1P-LIKE PROTEIN 14A Encodes a Bet1/Sft1-like | Yes | tRNA-Gly(CCC) | No | - |
| | | | GO:0015031 | protein transport | | | | | | |

| | | | | | | | | | | |
|---|---|---|---|---|---|---|---|---|---|---|
| | | | | | | SNARE protein which fully suppresses the temperature-sensitive growth defect in sft1-1 yeast cells; however, it cannot support the deletion of the yeast BET1 gene | | | | |
| VIT_14s0066g02600 | XP_019080906.1 | Protein arginine methyltransferase NDUFAF7 homolog, mitochondrial isoform | GO:0035243<br>GO:0032981 | protein-arginine omega-N symmetric methyltransferase activity<br>mitochondrial respiratory chain complex I assembly | AT3G28700 (75% AA ID, 2.5E-142 E-val) | NADH dehydrogenase ubiquinone complex I, assembly factor-like protein (DUF185) | No | - | No | - |
| VIT_13s0064g00200 | XP_002274669.1 | Uncharacterized protein LOC100267159 | No GO terms | No GO terms | AT4G19390 (59.2% AA ID, 3.3E-25 E-val) | Uncharacterized protein family (UPF0114) | No | - | Yes | EDTA-exudation (184850610) |
| VIT_17s0000g06990 | XP_019081977.1 | NAC transcription factor 32-like | GO:0003677<br>GO:0003735<br>GO:0006355<br>GO:0006412 | DNA binding<br>structural constituent of ribosome<br>regulation of transcription, DNA-templated<br>translation | AT1G33060 (48.9% AA ID, 4.6E-7 E-val) | ANAC014, NAC 014, NAC014 | No | - | No | - |
| VIT_18s0001g09050 | XP_010664507.1 | GPN-loop GTPase 2 | GO:0003924 | GTPase activity | AT4G12790 | GPN3<br>GPN GTPase | No | - | Yes | EDTA-exudation |

| | | | | | | | | | | |
|---|---|---|---|---|---|---|---|---|---|---|
| | | | GO:0005525 | GTP binding | (51.8% AA ID, 8.9E-26 E-val) | involved in selective nuclear import of RNA polymerase II. | | | | (18485061) |
| VIT_00s0322g00020 | XP_010646812.1 | Heptahelical transmembrane protein 4 | GO:0038023<br>GO:0009725 | signalling receptor activity<br>response to hormone | AT2G40710 (86% AA ID, 5.2E-55 E-val) | hemolysin-III related integral membrane protein | No | - | Yes | LMPC-exudation, EDTA-exudation (18485061) |
| VIT_04s0023g03700 | XP_002263940.1 | Uncharacterized protein LOC100267639 | No GO terms | No GO terms | AT4G35980 (77.6 AA ID, 1.2E-38 E-val) | hypothetical protein | No | - | Yes | LMPC-exudation (18485061) |
| VIT_07s0005g02210 | XP_010652305.1 | Uncharacterized protein LOC100257191 | GO:0005515 | protein binding | AT4G01860 (59.2% AA ID, 4.5E-97 E-val) | Transducin family protein / WD-40 repeat family protein | Yes | tRNA-Phe(GAA) | No | - |
| VIT_05s0077g01490 | XR_002030041.1 | Uncharacterized protein LOC104879211 | GO:0003676 | nucleic acid binding | AT5G26940 (40% AA ID, 6.0E-5 E-val) | DEFECTIVE IN POLLEN ORGANELLE DNA DEGRADATION1, DPD1 Encodes a Mg2+-dependent DNA exonuclease that degrades organelle DNA during Arabidopsis pollen development | No | - | No | - |
| VIT_19s0177g00220 | XP_002281637.1 | Electron transfer flavoprotein subunit alpha, mitochondrial | GO:0005507<br>GO:0009055<br>GO:0050660 | copper ion binding<br>electron transfer activity<br>flavin adenine dinucleotide binding | AT1G50940 (75.7 AA ID, 3.8E-160 E-val) | ELECTRON TRANSFER FLAVOPROTEIN ALPHA, ETFALPHA Encodes the electron transfer flavoprotein ETF alpha, a putative | No | - | No | - |

| | | | | | | | | | |
|---|---|---|---|---|---|---|---|---|---|
| | | | GO:0016491 | oxidoreductase activity | | subunit of the mitochondrial electron transfer flavoprotein complex (ETF beta is At5g43430.1) in Arabidopsis. Mutations of the ETF beta gene results in accelerated senescence and early death compared to wild-type during extended darkness. | | | |
| | | | GO:0033539 | fatty acid beta-oxidation using acyl-CoA dehydrogenase | | | | | |
| VIT_08s0007g03950 | XP_010654034.1 | Splicing factor U2af small subunit B | GO:0046872 | metal ion binding | AT1G30480 (34.1% AA ID, 0.07 E-val) | DNA-DAMAGE-REPAIR/TOLERATION PROTEIN 111, DRT111, REQUIRED FOR SNC4-1D 2, RSN2, SFPS, SPLICING FACTOR FOR PHYTOCHROME SIGNALING recombination and DNA-damage resistance protein | No | No | - |
| | | | GO:0030628 | pre-mRNA 3'-splice site binding | | | | | |
| | | | GO:0000398 | mRNA splicing, via spliceosome | | | | | |
