## Supplemental Table5 for "Tissue and regional expression patterns of dicistronic tRNA-mRNA transcripts in grapevine (*Vitis vinifera*) and their evolutionary co-appearance with vasculature in land plants"

**Supplemental TableS5**: Pearson correlation of the expression of tRNA and gene against intergenic region. Significant correlations are highlighted in green

| Leaf | Correlattion analysis (Pearson) of the expression of: | | | |
|---|---|---|---|---|
| | Dicistronic tRNAs vs intergenic region | | Dicistronic genes vs intergenic region | |
| region | correlation coefficient | p-value | correlation coefficient | p-value |
| CG | 0.76 | 0.0006241 | 0.217 | 0.4187 |
| EE | 0.638 | 0.007714 | 0.275 | 0.302 |
| EV | 0.769 | 0.0004872 | 0.258 | 0.3337 |
| NG | 0.663 | 0.005119 | 0.49 | 0.05378 |
| SG | 0.765 | 0.0005456 | 0.317 | 0.2303 |
| WR | 0.779 | 0.0003724 | 0.243 | 0.364 |

| Berry | Correlattion analysis (Pearson) of the expression of: | | | |
|---|---|---|---|---|
| | Dicistronic tRNAs vs intergenic region | | Dicistronic genes vs intergenic region | |
| region | correlation coefficient | p-value | correlation coefficient | p-value |
| CG | 0.666 | 0.07122 | -0.467 | 0.2432 |
| EE | 0.591 | 0.1224 | -0.445 | 0.2686 |
| EV | 0.679 | 0.06355 | -0.556 | 0.1518 |
| NG | 0.376 | 0.3577 | -0.445 | 0.2682 |
| SG | 0.703 | 0.0515 | -0.281 | 0.5 |
| WR | 0.69 | 0.05807 | -0.286 | 0.4915 |
