## Supplemental Table6 for "Tissue and regional expression patterns of dicistronic tRNA-mRNA transcripts in grapevine (*Vitis vinifera*) and their evolutionary co-appearance with vasculature in land plants"

Supplemental Table S6: List of primers used for cDNA synthesis and RT-PCR

| Name | primer | Sequence | Comments | Tissue expressed | Use for: | | |
|---|---|---|---|---|---|---|---|
| | | | | | cDNA synthesis | RT-PCR | Sanger sequencing |
| tRNA$^{ProTGG}$_F | Forward | TGCGAGAGGTCCCGAGTTCGATT | PCR product is 172 bp | Leaf & berry | | | |
| Intergenic_tRNA$^{ProTGG}$-VIT_18s0001g09050_R | Reverse | CTGTTTCCTTGCCTGCCACC | | | | | |
| VIT_18s0001g09050_R | Reverse | TGCATCATTGGCAGGATCCA | | | | | |
| tRNA$^{ValCAC}$_F | Forward | CACTAGAGGTCCCCGGTTCGAA | PCR product is 376 bp | Leaf | | | |
| Intergenic_tRNA$^{ValCAC}$-VIT_15s0046g02860_R | Reverse | CCGCAAGACCCAGATGGGAA | | | | | |
| VIT_15s0046g02860_R | Reverse | CCACCCCCTTTGAAGCCACA | | | | | |
| VvEF1-a_F | Forward | GAACTGGGTGCTTGATAGGC | PCR product is 150 bp | Leaf & berry | | | |
| VvEF1-a_R | Reverse | AACCAAAATATCCGGAGTAAAAGA | | | | | |
| tRNA$^{GlyCCC}$_R | Reverse | ACTAGATGCGCTGGATGAGG | | Berry | | | |
| VIT_19s0177g00220_F | Forward | TGGGACTTTAGTGTGGCAAA | PCR product is 367 bp | | | | |
| Intergenic_tRNA$^{GlyCCC-}$VIT_19s0177g00220_R | Reverse | TTTATGGTCCCTTTCCATGC | | | | | |
