## Supplemental Table7 for "Tissue and regional expression patterns of dicistronic tRNA-mRNA transcripts in grapevine (*Vitis vinifera*) and their evolutionary co-appearance with vasculature in land plants"

Identification of dicistronic tRNA-mRNA candidates in land plants using publicly available RNA-s

| Species | Reference genome used for read alignment |
| --- | --- |
| *Oryza sativa ssp. japonica* | *GCA_000005425.2 Build 4.0* |
| *Brachypodium distachyon* | *GCA_000005505.4 Brachypodium_distachyon_v3.0* |

| | |
|---|---|
| *Arabidopsis thaliana* | *GCA_000001735.2 TAIR10.1* |
| *Vitis vinifera* | |

*GCA_000003745.2*

| | |
|---|---|
| *Azolla Filiculoides* | *Azolla_asm_v1.1* |
| *Salvinia cucullata* | *Salvinia_asm_v1.2* |
| *Selaginella moellendorfii* | |

| | |
|---|---|
| | *GCA_000143415.2 v1.0* |

| | |
|---|---|
| *Physcomitrella patens* | *GCA_000002425.2 Phypa V3* |
| *Marchantia polymorpha* | GCA_003032435.1 Marchanta_polymorpha_v1 |

seq datasets from the NCBI-SRA repository

| SRA Project ID | SRA Accession (million reads) |
| --- | --- |
| PRJNA403975 | SRR6200382(61.9) SRR6200383(69.9) SRR6200376(57.3) |
| PRJNA301554 | SRR2931067(31.2) SRR2931068(28.9) SRR2931069(34.2) |
| PRJNA350792 | SRR4457249(12.6), SRR4457252(41.2), SRR4457253(31.0), SRR4457254(35.5) |
| PRJEB8082 | ERR748773(123.1), ERR748774(124.1), ERR748775(165.7), ERR748776(159.1), ERR748777(111.6) |
| PRJNA293888 | SRR3170524(54.1), SRR3170525(36.6), SRR3170526(42.4) |
| PRJNA371381 | SRR5270277(85.9), SRR5270278(101.1), SRR5270279(76.1) |
| PRJNA383406 | SRR5456708(25.6), SRR5456709(29.9), SRR5456710(28.9) |

| | |
|---|---|
| PRJNA271927 | SRR1773557(32.0),<br>SRR1773558(31.8),<br>SRR1773559(35.2)<br><br>SRR1773560(31.0)<br>SRR1773561(50.1)<br>SRR1773562(30.8)<br>SRR1773574(31.0)<br>SRR1773575(26.5)<br>SRR1773576(27.2) |
| PRJNA241198 | SRR1192255(56.0)<br>SRR1192256(61.0)<br>SRR1192257(67.1) |
| PRJNA246731 | SRR1283943(91.3)<br>SRR1283944(90.9)<br>SRR1283945(168.1) |
| PRJNA318918 | SRR3403548(31.8)<br>SRR3403549(31.5)<br>SRR3403550(31.1)<br>SRR3403551(31.0),<br>SRR3403552(31.5),<br>SRR3403553(31.2)<br><br>SRR3403554(30.6)<br>SRR3403555(33.0)<br>SRR3403556(31.5) |
| PRJNA311966 | SRR3167554 (44.1)<br>SRR3167555 (44.1)<br>SRR3167556 (38.2)<br><br>SRR3167560 (43.5)<br>SRR3167561 (37.7)<br>SRR3167562 (80.7)<br>SRR3167566 (63.1)<br>SRR3167567 (75.9)<br>SRR3167568 (67.1) |

| | |
|---|---|
| | SRR3167572 (68)<br>SRR3167573 (100.8)<br>SRR3167574 (61.3) |
| PRJNA383160 | SRR5453923 (15)<br>SRR5453924 (15.2)<br>SRR5453925 (19.6)<br>SRR5453929 (17)<br>SRR5453930 (20.2)<br>SRR5453931 (18.8) |
| PRJNA342391 | SRR4212894 (36.3)<br>SRR4212895 (47.3)<br>SRR4212896 (44.2) |
| PRJNA260535 | SRR1748366 (34.3)<br>SRR1748367 (34.5)<br>SRR1748368 (36.8)<br>SRR1748369 (35.1)<br>SRR1748370 (38)<br>SRR1748371 (30.3)<br><br>SRR1748339 (31.4)<br>SRR1748340 (33.7)<br>SRR1748341 (35)<br>SRR1748342 (27.9)<br>SRR1748343 (34.4)<br>SRR1748344 (29.3)<br>SRR1748345 (31)<br>SRR1748346 (36.9)<br>SRR1748347 (34.5) |
| PRJNA307079 | SRR3056884 (24.5)<br>SRR3056885 (24.9)<br>SRR3056886 (24.8)<br>SRR3056887 (23.9);<br>SRR3056888 (24.5);<br>SRR3056889 (21) |
| | SRR5534893 (50.2)<br>SRR5534894 (31.9)<br>SRR5534895 (33.4) |

| | |
|---|---|
| PRJNA386484 | SRR5534899 (49.5);<br>SRR5534900 (76.8);<br>SRR5534901 (32.2)<br><br><br><br>SRR5534905 (33.2)<br>SRR5534906 (30.2)<br>SRR5534907 (34.4) |
| PRJNA433817 | SRR6706481 (26)<br>SRR6706482 (25)<br>SRR6706484 (27.9) |
| PRJNA268857 | SRX807992 (35)<br>SRX807991 (32)<br>SRX807990 (24) |
| PRJNA430527 | SRR6480278 (45)<br>SRR6480271 (48)<br>SRR6480274 (39)<br>SRR6480273 (44)<br>SRR6480279 (46)<br>SRR6480280 (50) |
| PRJNA264391 | SRR1618558 (31)<br>SRR1618556 (30)<br>SRR1618551 (32) |
| PRJNA430459 | SRR6478600 (52)<br>SRR6478601 (40)<br>SRR6478602 (43)<br>SRR6478597 (36)<br>SRR6478598 (39)<br>SRR6478599 (49) |
| PRJEB29012 | ERR2820935 (33)<br>ERR2820936 (31)<br>ERR2820937 (34) |
| | SRR1740446 (36)<br>SRR1740447 (25)<br>SRR1740448 (38) |

| | |
|---|---|
| PRJNA271595 | SRR1740449 (29)<br>SRR1740450 (21)<br>SRR1740451 (28) |

| | |
|---|---|
| PRJNA417704 | SRR6279743 (31)<br>SRR6279744 (54)<br>SRR6279745 (29)<br>SRR6279746 (39) |
| PRJNA397394 | SRR5905095 16),<br>SRR5905096 (18)<br>SRR5905104 (16) |
| PRJNA294412 | SRR2225590 (31)<br>SRR2225591 (39)<br>SRR2225592 (31)<br>SRR2225593 (33) |
| PRJNA259146 | SRR1588451 (33)<br>SRR1588458 (48)<br>SRR1588464 (32) |
| PRJNA259147 | SRR1588577 (30)<br>SRR1588587 (34)<br>SRR1588592 (32) |
| PRJDB4420 | DRX045359 (14)<br>DRX045358 (14)<br>DRX045357 (19)<br>DRX045356 (15)<br>DRX045355 (17)<br>DRX045354 (14)<br>DRX045353 (16)<br>DRX045352 (15)<br>DRX045351 (18) |
| PRJNA433456 | SRX3661970 (38)<br>SRX3661969 (39)<br>SRX3661968 (36) |

| Tissue |
| --- |
| ecotype:Oryza sativa Japonica Group<br>tissue:root<br>no treatment |
| ecotype:Oryza sativa Japonica Group<br>tissue: leaf<br>no treatment |
| treatment:4 day after heading<br>treatment:12 day after heading |
| ecotype:Oryza sativa Japonica Group<br>tissue: leaf<br>no treatment |
| ecotype: Bd21-3<br>tissue: leaf<br>no treatment |
| ecotype: Bd21-3<br>tissue: leaf<br>non-vernalization |
| ecotype: Brachypodium distachyon<br>tissue: root<br>no treatment |

| |
|---|
| ecotype:col-1<br>tissue:root<br>no treatment<br><br>ecotype:col-1<br>tissue:shoot<br>no treatment<br>ecotype:Ped<br>tissue:root<br>no treatment |
| ecotype:col-1<br>tissue:leaves<br>no treatment |
| ecotype:col-1<br>tissue:leaves<br>no treatment |
| ecotype:col-1<br>tissue:leaves<br>heat 0 hour control<br>ecotype:col-1<br>tissue:leaves<br>heat 4 hour control<br><br>ecotype:col-1<br>tissue:leaves<br>heat 8 hour control |
| 2013 28 days after verison - wholle berry<br><br>2013 35 days after verison - whole berry<br><br>2014 28 days after verison - whole berry |

| |
|---|
| 2014 35 days after verison - whole berry |
| Non-inoculated non-wounded leaves<br><br>Non-inoculated non-wounded leaves |
| low nitrate 0 hpt -root tips |
| Sauvigon Banc berry skins at 24 hr brix<br><br>Sauvigon Banc berry skins at 24 hr brix<br><br>Sauvigon Banc berry skins at 22 hr brix<br><br>Sauvigon Banc berry skins at 24 hr brix<br><br>Sauvigon Banc berry skins at 26 hr brix |
| Vitis vinifera cv Prieto Picudo flower bud<br><br>Vitis vinifera cv Prieto Picudo flower bud |
| Vitis vinifera (cv. Sauvignon blanc) berry, light treatment - contr |

| |
|---|
| Vitis vinifera (cv. Sauvignon blanc) berry, light treatment - contr<br><br>Vitis vinifera (cv. Sauvignon blanc) berry, light treatment - contr |
| Vitis vinifera (cv. Pinot Noir) leaf, treatment - control<br><br>Vitis vinifera (cv. Pinot Noir) berry skins |
| Azolla Filiculoides whole plant<br><br>Azolla Filiculoides whole plant |
| Root tip sample of Azolla filiculoides mock treated |
| Salvinia cucullata floating leaves<br><br>Salvinia cucullata submerged leaves |
| vascular leaf |
| meristematic zone (MZ1-3) |

| |
|---|
| root tips (EDZ1-3) |

| |
|---|
| ecotype:<br>tissue: protonema<br>no treatment |
| ecotype:<br>tissue: Protonema<br>no treatment |
| ecotype:<br>tissue: protonema<br>no treatment<br><br>Physcomitrella patens<br><br>Physcomitrella patens |
| archegoniophore<br><br>antheridiophore<br><br>antheridiophore |
| vegetative thallus |

| DiRT identifed tRNA-mRNA dicistronic candidate | Interval (Distance in bp between tRNA and PC( |
|---|---|
| chr10.trna30-GlyGCC_Os10g0565300 | 1156 |
| chr10.trna32-MetCAT_Os10g0563500 | 150 |
| chr1.trna28-ThrAGT_Os01g0767000 | 3709 |
| chr1.trna58-PheGAA_Os01g0959800 | 1641 |
| chr1.trna67-LeuTAG_Os01g0861700 | 338 |
| chr1.trna67-LeuTAG_Os01g0861900 | 173 |
| chr1.trna82-TyrGTA_Os01g0759800 | 18 |
| chr3.trna22-TyrGTA_Os03g0616500 | 135 |
| chr4.trna41-HisGTG_Os04g0472700 | 2453 |
| chr5.trna45-ValCAC_Os05g0414100 | 2424 |
| chr1.trna67-LeuTAG_Os01g0861700 | 338 |
| chr3.trna22-TyrGTA_Os03g0616500 | 135 |
| chr10.trna17-IleAAT_Os10g0467600 | 24 |
| chr11.trna32-AlaAGC_Os11g0150100 | 210 |
| chr1.trna42-LysCTT_Os01g0841200 | 453 |
| chr1.trna82-TyrGTA_Os01g0759800 | 18 |
| chr11.trna32-AlaAGC_Os11g0150100 | 210 |
| chr11.trna32-AlaAGC_Os11g0150200 | 1563 |
| chr1.trna28-ThrAGT_Os01g0767000 | 3709 |
| chr1.trna42-LysCTT_Os01g0841200 | 453 |
| chr1.trna57-ArgTCT_Os01g0960400 | 834 |
| chr1.trna67-LeuTAG_Os01g0861900 | 173 |
| chr1.trna82-TyrGTA_Os01g0759800 | 18 |
| chr3.trna12-AlaAGC_Os03g0343300 | 149 |
| chr3.trna22-TyrGTA_Os03g0616500 | 135 |
| chr3.trna32-GlnCTG_Os03g0769000 | 502 |
| chr5.trna45-ValCAC_Os05g0414100 | 2424 |
| chr7.trna29-AlaCGC_Os07g0686400 | 690 |
| chr7.trna29-AlaCGC_Os07g0686500 | 2922 |
| BRADI_3g27341v3_chr3.tRNA98-LeuCAA | 15 |
| BRADI_3g27341v3_chr3.tRNA98-LeuCAA | 15 |
| BRADI_3g53300v3_chr3.tRNA59-GlnCTG | 192 |

| | | |
|---|---|---|
| AT2G18450_chr2.trna84-Undet??? | | 1831 |
| AT5G22320_tRNA-Gln-CTG-2-2 | | 25 |
| AT4G14410_tRNA-Thr-AGT-1-2 | | 56 |
| AT2G18450_chr2.trna84-Undet??? | | 1831 |
| AT3G20362_tRNA-Tyr-GTA-14-1 | | 57 |
| AT3G20362_tRNA-Tyr-GTA-14-1 | | 57 |
| AT3G09600_tRNA-Ser-TGA-4-1 | | 461 |
| AT1G15080_tRNA-Asn-GTT-8-1 | | 12 |
| AT1G28710_tRNA-Pro-AGG-5-1 | | 124 |
| AT4G36210_tRNA-Glu-CTC-4-4 | | 121 |
| AT1G28765_tRNA-Pro-TGG-2-20 | | 2 |
| AT1G54680_tRNA-Ala-TGC-1-5 | | 11 |
| AT2G07662_tRNA-Asn-GTT-7-1 | | 561 |
| tRNA-Gly-TCC-1-6_VIT_13s0064g00200 | 82 | |
| tRNA-Phe-GAA-1-4_VIT_07s0005g02200 | 89 | |
| tRNA-Pro-TGG-2-9_VIT_18s0001g09050 | 133 | |
| tRNA-Val-AAC-1-2_VIT_16s0100g00640 | 154 | |
| tRNA-Ser-CGA-1-2_VIT_06s0004g07100 | 222 | |
| tRNA-Gly-CCC-1-3_VIT_19s0177g00220 | 469 | |
| tRNA-Phe-GAA-1-4_VIT_07s0005g02200 | 89 | |
| tRNA-Pro-TGG-2-9_VIT_18s0001g09050 | 133 | |

| | |
|---|---|
| tRNA-Ala-TGC-3-1_VIT_07s0104g01010 | 98 |
| tRNA-Ile-AAT-3-1_VIT_04s0023g03700 | 105 |
| tRNA-Phe-GAA-1-4_VIT_07s0005g02200 | 89 |
| tRNA-Pro-TGG-2-9_VIT_18s0001g09050 | 133 |
| tRNA-Asn-GTT-4-1_VIT_18s0072g00300 | 223 |
| tRNA-Gly-TCC-1-6_VIT_13s0064g00200 | 82 |
| tRNA-Arg-CCG-3-1_VIT_04s0008g05550 | 171 |
| tRNA-Pro-TGG-2-9_VIT_18s0001g09050 | 133 |
| tRNA-Pro-TGG-2-9_VIT_18s0001g09050 | 133 |
| tRNA-His-GTG-8-1_VIT_17s0000g06990 | 13 |
| tRNA-Pro-TGG-2-9_VIT_18s0001g09050 | 133 |
| tRNA-Pro-TGG-2-9_VIT_18s0001g09050 | 133 |
| tRNA-Pro-TGG-2-9_VIT_18s0001g09050 | 133 |
| tRNA-Phe-GAA-1-4_VIT_07s0005g02200 | 89 |
| tRNA-Pro-TGG-2-9_VIT_18s0001g09050 | 133 |
| tRNA-Gln-CTG-4-1_VIT_02s0025g01690 | 64 |
| tRNA-Asp-GTC-1-10_VIT_17s0000g03980 | 179 |
| tRNA-Pro-CGG-2-2_VIT_14s0060g01370 | 129 |
| tRNA-Thr-AGT-2-1_VIT_05s0077g01490 | 562 |
| tRNA-Pro-TGG-2-9_VIT_18s0001g09050 | 133 |

| | |
|---|---|
| tRNA-Gly-TCC-1-6_VIT_13s0064g00200 | 82 |
| tRNA-Ile-AAT-3-1_VIT_04s0023g03700 | 105 |
| tRNA-Pro-TGG-2-9_VIT_18s0001g09050 | 133 |
| tRNA-Thr-AGT-1-4_VIT_00s0322g00020 | 82 |
| tRNA-Pro-TGG-2-9_VIT_18s0001g09050 | 133 |
| tRNA-Gln-CTG-4-1_VIT_02s0025g01690 | 64 |
| tRNA-Asn-GTT-2-2_VIT_18s0001g12620 | -700 |
| tRNA-Ala-TGC-3-1_VIT_07s0104g01010 | 98 |
| tRNA-Phe-GAA-1-4_VIT_07s0005g02200 | 89 |
| tRNA-Pro-TGG-2-9_VIT_18s0001g09050 | 133 |
| Azfi_s0030-5TrpCCA_Azfi_s0030.g024311 | 99 |
| Azfi_s0030-5TrpCCA_Azfi_s0030.g024311 | 99 |
| Azfi_s0030-5TrpCCA_Azfi_s0030.g024311 | 99 |
| Azfi_s0804-1ArgACG_Azfi_s0804.g087566 | 251 |
| Sacu_v1.1_s0003-33LeuCAG_Sacu_v1.1_s0003.g0020 | 174 |
| Sacu_v1.1_s0126-1LeuCAG_Sacu_v1.1_s0126.g02173 | 1893 |
| intergenic_GL377628.tRNA-LeuCAA_SELMODRAFT_23 | 2965 |
| intergenic_GL377614.tRNA TyrGUA_SELMODRAFT_42 | 290 |
| intergenic_GL377569.tRNA-ArgUCU_SELMODRAFT_27 | 10 |
| intergenic_GL377576.tRNA-PheGAA_SELMODRAFT_9 | 112 |
| intergenic_GL377573.tRNA-TyrGUA_SELMODRAFT_67 | 140 |
| intergenic_GL377565.tRNA12-ArgACG_SELMODRAFT_ | 1035 |
| intergenic_GL377565.tRNA18-AsnGTT_SELMODRAFT_ | 87 |
| intergenic_GL377565.tRNA21-ValTAC_SELMODRAFT_ | 186 |
| intergenic_GL377565.tRNA29-UndetNNN_SELMODRA | 405 |

| | |
|---|---|
| intergenic_GL377565.tRNA30-PheGAA_SELMODRAFT_ | 148 |
| intergenic_GL377569.tRNA33-CysGCA_SELMODRAFT_ | 85 |
| intergenic_GL377569.tRNA36-ArgTCT_SELMODRAFT_ | 1 |
| intergenic_GL377573.tRNA7-ValGAC_SELMODRAFT_2 | 136 |
| intergenic_GL377577.tRNA14-AsnGTT_SELMODRAFT_ | 87 |
| intergenic_GL377577.tRNA17-ValTAC_SELMODRAFT_ | 189 |
| intergenic_GL377589.tRNA20-ValAAC_SELMODRAFT_ | 166 |
| intergenic_GL377594.tRNA9-UndetNNN_SELMODRAF | 66 |
| intergenic_GL377604.tRNA4-IleGAT_SELMODRAFT_10 | 2251 |
| intergenic_GL377604.tRNA4-IleGAT_SELMODRAFT_25 | 823 |
| intergenic_GL377620.tRNA1-AlaTGC_SELMODRAFT_1 | 55 |
| intergenic_GL377628.tRNA8-LeuCAA_SELMODRAFT_2 | 2964 |
| intergenic_GL377629.tRNA3-ArgTCG_SELMODRAFT_4 | 45 |
| intergenic_GL377634.tRNA12-ValAAC_SELMODRAFT_ | 164 |
| intergenic_GL377639.tRNA1-IleGAT_SELMODRAFT_12 | 2224 |
| intergenic_GL377645.tRNA3-HisGTG_SELMODRAFT_4 | 109 |
| intergenic_GL377677.tRNA5-ProCGG_SELMODRAFT_2 | 204 |
| intergenic_GL377677.tRNA6-ProAGG_SELMODRAFT_2 | 55 |
| intergenic_GL377714.tRNA5-GluTTC_SELMODRAFT_2 | 261 |
| intergenic_GL377565.tRNA12-ArgACG_SELMODRAFT_ | 1035 |
| intergenic_GL377565.tRNA21-ValTAC_SELMODRAFT_ | 186 |
| intergenic_GL377569.tRNA10-GluTTC_SELMODRAFT_ | 269 |
| intergenic_GL377569.tRNA33-CysGCA_SELMODRAFT_ | 85 |
| intergenic_GL377569.tRNA36-ArgTCT_SELMODRAFT_ | 1 |
| intergenic_GL377569.tRNA9-GluCTC_SELMODRAFT_1 | 1746 |
| intergenic_GL377576.tRNA7-AlaAGC_SELMODRAFT_6 | 627 |
| intergenic_GL377577.tRNA17-ValTAC_SELMODRAFT_ | 189 |
| intergenic_GL377581.tRNA7-MetCAT_SELMODRAFT_ | 599 |
| intergenic_GL377589.tRNA20-ValAAC_SELMODRAFT_ | 166 |
| intergenic_GL377595.tRNA3-LeuTAA_SELMODRAFT_2 | 46 |
| intergenic_GL377596.tRNA3-HisGTG_SELMODRAFT_1 | 320 |
| intergenic_GL377607.tRNA2-ArgTCT_SELMODRAFT_5 | 123 |
| intergenic_GL377620.tRNA1-AlaTGC_SELMODRAFT_1 | 55 |
| intergenic_GL377622.tRNA9-ArgACG_SELMODRAFT_4 | 71 |
| intergenic_GL377626.tRNA10-LysTTT_SELMODRAFT_1 | 399 |
| intergenic_GL377632.tRNA3-ArgTCT_SELMODRAFT_4 | 123 |
| intergenic_GL377634.tRNA12-ValAAC_SELMODRAFT_ | 164 |
| intergenic_GL377675.tRNA1-PheGAA_SELMODRAFT_ | 111 |
| intergenic_GL377677.tRNA5-ProCGG_SELMODRAFT_2 | 204 |
| intergenic_GL377677.tRNA6-ProAGG_SELMODRAFT_2 | 55 |
| intergenic_GL377692.tRNA3-GluCTC_SELMODRAFT_1 | 1747 |
| intergenic_GL377714.tRNA5-GluTTC_SELMODRAFT_2 | 261 |

G)
