## Supplemental Table8 for "Tissue and regional expression patterns of dicistronic tRNA-mRNA transcripts in grapevine (*Vitis vinifera*) and their evolutionary co-appearance with vasculature in land plants"

Supplementary Table S8: Lignin related genes

| | *A.thaliana* | *V. vinifera* | *O. sativa* | *B.distachyon* | *A. filiculoides* | *S. cucullata* | *S.moellendorffii* | *P. patens* | *M. polymorpha* |
|---|---|---|---|---|---|---|---|---|---|
| **PAL1** | At2g37040 | GSVIVT01024292001 | LOC_Os02g41650 | Bradi3g49280 | Azfi_s0009.g011807 | Sacu_v1.1_s0091.g018764 | 424403 | Pp3c14_11870V3.1.p | Mapoly0070s0071/Mp4g14110 |
| **PAL2** | At3g53260 | ? | LOC_Os04g43800 | Bradi5g15830 | Azfi_s0063.g035303 | Sacu_v1.1_s0253.g026791 | n.d. | Pp3c1_18940V3.1.p | Mapoly0070s0068/Mp4g14140 |
| **PAL4** | At3g10340 | GSVIVT01025214001 | LOC_Os02g41680 | Bradi3g49260 | Azfi_s0096.g043679 | Sacu_v1.1_s0017.g007129 | 404263 | Pp3c1_18830V3.1.p | Mapoly0014s0211/Mp1g10150 |
| **C4H** | At2g30490 | GSVIVT01024554001 | LOC_Os05g25640 | Bradi2g53470 | Azfi_s0333.g065437 | Sacu_v1.1_s0039.g012203 | 175973 | Pp3c4_21680V3.1.p | Mapoly0163s0018/Mp6g00020 |
| **4CL1** | At1g51680 | GSVIVT01029182001 | LOC_Os02g08100 | Bradi3g05750 | Azfi_s0114.g046013 | Sacu_v1.1_s0149.g023264 | 171251 | Pp3c18_6360V3.1.p | Mapoly0197s0014/Mp5g01200 |
| **4CL2** | At3g21240 | GSVIVT01029183001 | LOC_Os06g44620 | Bradi3g52350 | Azfi_s0013.g013344 | n.d. | 177393 | Pp3c19_13170V3.1.p | n.d. |
| **HCT** | At5g48930 | GSVIVT01016053001 | LOC_Os04g42250 | Bradi5g14720 | Azfi_s0005.g009338 | Sacu_v1.1_s0010.g004618 | 152997 | Pp3c2_29140V3.1.p | Mapoly0003s0277/Mp7g12690 |
| **C3H1** | At2g40890 | GSVIVT01025800001 | LOC_Os05g41440 | Bradi2g21300 | Azfi_s0355.g066730 | Sacu_v1.1_s0001.g000031 | 271465 | Pp3c22_19010V3.1.p | Mapoly0037s0087/Mp3g11100 |
| **CSE** | At1g52760 | GSVIVT01017214001 | LOC_Os02g11720 | Bradi1g24490 | Azfi_s0045.g030088 | Sacu_v1.1_s0674.g027619 | 113971 | Pp3c19_14430V3.1.p | Mapoly0032s0047/Mp5g13540 |
| **CCoAOMT1** | At4g34050 | GSVIVT01022100001 | LOC_Os06g06980 | Bradi1g48370 | Azfi_s0002.g001302 | Sacu_v1.1_s0016.g006795 | 80209 | Pp3c23_5530V3.1.p | Mapoly0035s0134/Mp6g03550 |
| **CCoAOMT7** | At4g26220 | GSVIVT01015245001 | LOC_Os08g38910 | Bradi4g33340 | Azfi_s0167.g054536 | ? | 271191 | ? | Mapoly0099s0056/Mp7g01830 |
| **CCR1** | At1g15950 | GSVIVT01034241001 | LOC_Os08g34280 | Bradi3g36887 | Azfi_s3139.g114683 | Sacu_v1.1_s0016.g006953 | 271114 | Pp3c7_17190V3.1.p | ? |
| **F5H1** | At4g36220 | GSVIVT01024186001 | LOC_Os10g36848 | Bradi3g30590 | n.d. | n.d. | n.d. | n.d. | n.d. |
| **COMT** | At5g54160 | GSVIVT01008854001 | LOC_Os08g06100 | Bradi3g16530 | Azfi_s0019.g015239 | Sacu_v1.1_s0056.g014618 | 438615 | Pp3c12_5860V3.1.p | Mapoly0337s0001/Mp2g07360 |
| **CAD3** | At4g34230 | GSVIVT01003150001 | LOC_Os02g09490 | Bradi3g06480 | Azfi_s0001.g000474 | Sacu_v1.1_s0209.g025876 | 444096 | ? | ? |
| **CAD6** | At4g37970 | GSVIVT01006303001 | LOC_Os10g29470 | Bradi3g17920 | Azfi_s0021.g015839 | Sacu_v1.1_s0027.g009722 | 230239 | Pp3c1_39700V3.1.p | ? |
| **LAC4** | At2g38080 | GSVIVT01034003001 | LOC_Os11g48060 | Bradi1g74320 | Azfi_s0374.g067219 | Sacu_v1.1_s0058.g014969 | 165365 | Pp3c15_19050V3.1.p | n.d. |
| **LAC11** | At5g03260 | GSVIVT01025694001 | LOC_Os01g44330 | Bradi2g54680 | Azfi_s1432.g103020 | Sacu_v1.1_s0039.g012147 | 404075 | Pp3c1_13930V3.1.p | Mapoly0049s0002/Mp3g20310 |
| **LAC17** | At5g60020 | GSVIVT01025046001 | LOC_Os01g62480 | Bradi1g24880 | Azfi_s0402.g068306 | Sacu_v1.1_s0031.g010585 | 95740 | Pp3c6_3290V3.1.p | n.d. |

n.d. not detected

? unequivocal ortholog assignment not possible
