## Supplemental Table9 for "Tissue and regional expression patterns of dicistronic tRNA-mRNA transcripts in grapevine (*Vitis vinifera*) and their evolutionary co-appearance with vasculature in land plants"

**Supplementary Table S10: Secondary cell wall celluolose related genes**

| | *A. thaliana* | *V. vinifera* | *O. sativa* | *B. distachyon* | *A. filiculoides* | *S. cucullata* | *S.moellendorffii* | *P. patens* | *M. polymorpha* |
|---|---|---|---|---|---|---|---|---|---|
| MYB46 | At5g12870 | n.d. | n.d. | n.d. | n.d. | n.d. | n.d. | n.d. | n.d. |
| CesA4/IRX5 | At5g44030 | GSVIVT01028402001 | LOC_Os10g32980.1 | Bradi3g28350 | Azfi_s0007.g010884 | ? | ? | ? | n.d. |
| CesA7/IRX3 | At5g17420 | GSVIVT01023643001 | LOC_Os09g25490.1 | Bradi4g30540 | Azfi_s0059.g034631 | Sacu_v1.1_s0085.g018303 | 163575 | Pp3c9_11990V3.1.p | n.d. |
| CesA8/IRX1 | At4g18780 | GSVIVT01021248001 | LOC_Os01g54620.1 | Bradi2g49912 | Azfi_s0230.g059145 | ? | ? | ? | n.d. |
| KORRIGAN | At5g49720 | GSVIVT01023102001 | LOC_Os03g52630.1 | Bradi1g09460 | Azfi_s0006.g009785 | Sacu_v1.1_s0149.g023273 | 75214 | Pp3c3_27980V3.1.p | Mapoly0061s0043/Mp1g24780 |
| COBL4 | At5g15630 | GSVIVT01036565001 | LOC_Os03g30250.1 | Bradi1g59880 | Azfi_s0018.g014792 | Sacu_v1.1_s0125.g021681 | 271954 | Pp3c12_4550V3.1.p | Mapoly0057s0007/Mp7g06600 |
| TED6 | At1g43790 | n.d. | n.d. | Bradi4g45531 | n.d. | n.d. | n.d. | n.d. | n.d. |

n.d. not detected

? unequivocal ortholog assignment not possible
