## Supplemental Table10 for "Tissue and regional expression patterns of dicistronic tRNA-mRNA transcripts in grapevine (*Vitis vinifera*) and their evolutionary co-appearance with vasculature in land plants"

Supplementary table S11: Number of tRNA-mRNA identified for each tRNA isodecoders in independent RNA-seq datasets (NCBI-SRA) in vascular and non-vascular land plants

| **Amino acid** | **Anticodon** | ***Selaginella moellendorfii*** | ***Salvinia cucullata*** | ***Azolla Filiculoides*** | ***Vitis vinifera*** | ***Arabidopsis thaliana*** | ***Brachypodium distachyon*** | ***Oryza sativa*** | ***Marchantia polymorpha*** | ***Physcomitrella patens*** |
|---|---|---|---|---|---|---|---|---|---|---|
| Ala | AGC | 1 | | | | | | *4* | | |
| | CGC | | | | | | | *2* | | |
| | TGC | 2 | | | 2 | *1* | | | | |
| Arg | ACG | 3 | | 1 | | | | | | |
| | CCG | | | | 1 | | | | | |
| | TCG | 1 | | | | | | | | |
| | TCT | 4 | | | | | | *1* | | |
| | UCU | 1 | | | | | | | | |
| Asn | GTT | 2 | | | 2 | *2* | | | | |
| Asp | GTC | | | | 1 | | | | | |
| Cys | GCA | 2 | | | | | | | | |
| Gln | CTG | | | | 2 | *1* | *1* | *1* | | |
| Glu | CTC | 2 | | | | *1* | | | | |
| | TTC | 3 | | | | | | | | |
| Gly | CCC | | | | 1 | | | | | |
| | GCC | | | | | | | *1* | | |
| | TCC | | | | 4 | | | | | |
| His | GTG | 2 | | | 1 | | | *1* | | |
| Ile | AAT | | | | 2 | | | *1* | | |
| | GAT | 3 | | | | | | | | |
| Leu | CAA | 2 | | | | | *2* | | | |
| | CAG | | 2 | | | | | | | |
| | TAA | 1 | | | | | | | | |
| | TAG | | | | | | | *4* | | |

| | | | | | | | | |
|---|---|---|---|---|---|---|---|---|
| Lys | CTT | | | | | | | *2* |
| | TTT | 1 | | | | | | |
| Met | CAT | 1 | | | | | | *1* |
| Phe | GAA | 3 | | | 5 | | | *1* |
| Pro | AGG | 2 | | | | *1* | | |
| | CGG | 2 | | | 1 | | | |
| | TGG | | | | 13 | *1* | | |
| Ser | CGA | | | | 1 | | | |
| | TGA | | | | | *1* | | |
| Thr | AGT | | | | 2 | *1* | | *2* |
| Trp | CCA | | | 3 | | | | |
| Tyr | GTA | | | | | *2* | | *6* |
| | GUA | 2 | | | | | | |
| Val | AAC | 4 | | | 1 | | | |
| | CAC | | | | | | | *2* |
| | GAC | 1 | | | | | | |
| | TAC | 4 | | | | | | |
