## Supplemental material for "Tissue and regional expression patterns of dicistronic tRNA-mRNA transcripts in grapevine (*Vitis vinifera*) and their evolutionary co-appearance with vasculature in land plants"

Alignment of sequencing results from PCR product for two dicistronic tRNA-mRNA candidates to the expected PCR product.

```
Query: tRNA-Pro-TGG-2-9_Intergenic_region Query ID: lcl|Query_54328 Length:
171

>1_ProC_F_B09

Sequence ID: Query_54330 Length: 125

Range 1: 1 to 124


Score:224 bits(121), Expect:9e-64,

Identities:123/124(99%),  Gaps:0/124(0%), Strand: Plus/Plus


Query  48   TTTCTTTCGCTGGGTTTTGGTTTTACTTCACCATAAACCTCAAAAAAGCCCTCTTATGCT  107
            ||||||| ||||||||||||||||||||||||||||||||||||||||||||||||||||
Sbjct  1    TTTCTTTGGCTGGGTTTTGGTTTTACTTCACCATAAACCTCAAAAAAGCCCTCTTATGCT  60


Query  108  CTTCTGCAAATTTCATTTGTGTTATTGGTACTGAAACTCCGAGGCGGTGGCAGGCAAGGA  167
            ||||||||||||||||||||||||||||||||||||||||||||||||||||||||||||
Sbjct  61   CTTCTGCAAATTTCATTTGTGTTATTGGTACTGAAACTCCGAGGCGGTGGCAGGCAAGGA  120


Query  168  AACA  171
            ||||
Sbjct  121  AACA  124
```

Query: tRNA-Pro-TGG-2-9_Intergenic_region Query ID: lcl|Query_54328 Length: 171

>2_ProC_R_B10

Sequence ID: Query_54331 Length: 120

Range 1: 3 to 120

Score:213 bits(115), Expect:2e-60,

Identities:118/119(99%), Gaps:1/119(0%), Strand: Plus/Minus

```
Query  1    TGCGAGAGGTCCCGAGTTCGATTCTCGGAATGCCCCAATCTTTTTATTTTCTTTCGCTGG  60
            ||||||||||||||||||||||||||||||||||||||||||||||||||||||||||||
Sbjct  120  TGCGAGAGGTCCCGAGTTCGATTCTCGGAATGCCCCAATCTTTTTATTTTCTTTCGCTGG  61

Query  61   GTTTTGGTTTTACTTCACCATAAACCTCAAAAAAGCCCTCTTATGCTCTTCTGCAAATT  119
            ||||||||||||||||||||||||||||||||||||||||||||||||| |||||||||
Sbjct  60   GTTTTGGTTTTACTTCACCATAAACCTCAAAAAAGCCCTCTTATGCTCT-CTGCAAATT  3
```

Query: Val_intergenic_PCR_Product Query ID: lcl|Query_16750 Length: 376

>5_ValC_F_C01

Sequence ID: Query_16752 Length: 320

Range 1: 1 to 320

Score:586 bits(317), Expect:6e-172,

Identities:319/320(99%),  Gaps:0/320(0%), Strand: Plus/Plus

```
Query  56   ATTGCCAGAGTCTTCCATTTCTGTTGGGAGTCTCCCAGGGTCAGAGTATCAACGACACTC  115
            ||||||| ||||||||||||||||||||||||||||||||||||||||||||||||||||
Sbjct  1    ATTGCCACAGTCTTCCATTTCTGTTGGGAGTCTCCCAGGGTCAGAGTATCAACGACACTC  60

Query  116  AGTGCCACAATCTCATTTCCATTTCTGCTAGGAGTCTCCCAGATTCTCAATATCAACAAC  175
            ||||||||||||||||||||||||||||||||||||||||||||||||||||||||||||
Sbjct  61   AGTGCCACAATCTCATTTCCATTTCTGCTAGGAGTCTCCCAGATTCTCAATATCAACAAC  120

Query  176  ACTCACCCAGATTTTTAACATTTTCTCATCTGGATGTTCATCAATTAGTCAAACAATGCA  235
            ||||||||||||||||||||||||||||||||||||||||||||||||||||||||||||
Sbjct  121  ACTCACCCAGATTTTTAACATTTTCTCATCTGGATGTTCATCAATTAGTCAAACAATGCA  180

Query  236  GATTCAGCCACACCCACTTCCAAACTATAGTCTTAGGTCACCAATTTTCTCACCTTCGAC  295
            ||||||||||||||||||||||||||||||||||||||||||||||||||||||||||||
Sbjct  181  GATTCAGCCACACCCACTTCCAAACTATAGTCTTAGGTCACCAATTTTCTCACCTTCGAC  240

Query  296  CCATTTTCTCATACCTTTCTCATCATCATTCCAACCTACTAAACTTGTCTCTATTCCAAA  355
            ||||||||||||||||||||||||||||||||||||||||||||||||||||||||||||
Sbjct  241  CCATTTTCTCATACCTTTCTCATCATCATTCCAACCTACTAAACTTGTCTCTATTCCAAA  300
```

```
Query  356  TTTCCCATCTGGGTCTTGCG  375
            ||||||||||||||||||||
Sbjct  301  TTTCCCATCTGGGTCTTGCG  320
```

Query: Val_intergenic_PCR_Product Query ID: lcl|Query_16750 Length: 376

>6_ValC_R_C02

Sequence ID: Query_16753 Length: 307

Range 1: 1 to 307

Score:562 bits(304), Expect:1e-164,

Identities:306/307(99%),  Gaps:0/307(0%), Strand: Plus/Minus

```
Query  1    CACTAGAGGTCCCCGGTTCGAACCCGGGCTCAGACATTTGCATTTTTATTTTATTATTGC  60
            ||||||||||||||||||||||||||||||||||||||||||||||||||||||||||||
Sbjct  307  CACTAGAGGTCCCCGGTTCGAACCCGGGCTCAGACATTTGCATTTTTATTTTATTATTGC  248

Query  61   CAGAGTCTTCCATTTCTGTTGGGAGTCTCCCAGGGTCAGAGTATCAACGACACTCAGTGC  120
            || |||||||||||||||||||||||||||||||||||||||||||||||||||||||||
Sbjct  247  CACAGTCTTCCATTTCTGTTGGGAGTCTCCCAGGGTCAGAGTATCAACGACACTCAGTGC  188

Query  121  CACAATCTCATTTCCATTTCTGCTAGGAGTCTCCCAGATTCTCAATATCAACAACACTCA  180
            ||||||||||||||||||||||||||||||||||||||||||||||||||||||||||||
Sbjct  187  CACAATCTCATTTCCATTTCTGCTAGGAGTCTCCCAGATTCTCAATATCAACAACACTCA  128

Query  181  CCCAGATTTTTAACATTTTCTCATCTGGATGTTCATCAATTAGTCAAACAATGCAGATTC  240
            ||||||||||||||||||||||||||||||||||||||||||||||||||||||||||||
Sbjct  127  CCCAGATTTTTAACATTTTCTCATCTGGATGTTCATCAATTAGTCAAACAATGCAGATTC  68

Query  241  AGCCACACCCACTTCCAAACTATAGTCTTAGGTCACCAATTTTCTCACCTTCGACCCATT  300
            ||||||||||||||||||||||||||||||||||||||||||||||||||||||||||||
Sbjct  67   AGCCACACCCACTTCCAAACTATAGTCTTAGGTCACCAATTTTCTCACCTTCGACCCATT  8
```

```
Query  301  TTCTCAT  307
            |||||||
Sbjct  7    TTCTCAT  1
```
